# Pisces: A multi-modal data augmentation approach for drug combination synergy prediction

**DOI:** 10.1101/2022.11.21.517439

**Authors:** Hanwen Xu, Jiacheng Lin, Addie Woicik, Zixuan Liu, Jianzhu Ma, Sheng Zhang, Hoifung Poon, Liewei Wang, Sheng Wang

## Abstract

Drug combination therapy is promising for cancer treatment through simultaneously reducing resistance and improving efficacy. Machine learning approaches to drug combination response prediction can prioritize experiments and discover new combinations, but require lots of training data in order to fit the nonlinearity of synergistic effect. Here, we propose Pisces, a novel machine learning approach for drug combination synergy prediction. The key idea of Pisces is to augment the sparse drug combination dataset by creating multiple views for each drug combination based on its different modalities. We combined eight different modalities of a single drug to create 64 augmented views for a pair of drugs, effectively expanding the size of the original data 64 times. Pisces obtained state-of-the-art results on cell-line-based drug synergy prediction, xenograft-based drug synergy prediction, and drug-drug interaction prediction. By interpreting Pisces’s predictions using a genetic interaction network, we further identified a breast cancer drug-sensitive pathway from BRCA cell lines in GDSC. We validated this pathway on an independent TCGA-BRCA tumor dataset and found that patients with this pathway activated had substantially longer survival time. Collectively, Pisces effectively predicts drug synergy and drug-drug interactions through augmenting the original dataset 64 times, and can be broadly applied to various biological applications that involve a pair of drugs.

## Introduction

Drug combination therapy, which exploits two synergistic drugs whose combined effect is greater than the sum of each drug’s individual activity, is a promising treatment for cancer.^1–9^ A few successful drug combinations,^6,9–13^ such as lenvatinib and gefitinib for hepatocellular carcinoma (HCC),^10^ have delivered encouraging results in clinics. As a result, large-scale cancer pharmacology projects, including the Genomics of Drug Sensitivity in Cancer (GDSC) projects,^14^ Beat Acute Myeloid Leukemia project,^15^ national cancer institute ALMANAC project,^16^ and AstraZeneca’s DREAM challenge,^17^ have generated a large volume of genomics and pharmacological data for studying drug synergy, offering an unprecedented opportunity to discover new drug combinations and achieve precision medicine.^18^ Since it is expensive to experimentally profile all combinations, there is a pressing need to develop machine learning approaches for predicting drug combination response.^19–24^

Developing machine learning approaches for predicting drug combination response is inherently challenging. On the one hand, we need an expressive model, which requires lots of training data, to capture the nonlinearity of synergistic effect.^19–23^ On the other hand, we can only obtain limited training data due to the cubic growth of the drug combination space (i.e., two drugs and a cell line), which leads to the overfitting of an expressive model. Recently, data augmentation has transformed computer vision and natural language processing,^25–30^ and became an essential component in generative artificial intelligence.^31–36^ Data augmentation alleviates the overfitting of expressive machine learning models by creating multiple views of the same data instance and encouraging them to be similar to each other.^25^

Here we proposed Pisces, a novel machine learning model for drug combination synergy prediction. The key idea of Pisces is to augment the sparse drug combination dataset by creating multiple views for each drug combination based on different modalities. Our intuition is that different drug modalities are complementary to each other, and jointly considering them can enhance the prediction performance. Specifically, we consider eight drug modalities (**Fig. 1a**), including Simplified Molecular Input Line Entry System (SMILES),^37^ molecular graphs,^37^ three-dimensional molecular structures,^38^ drug targets,^39^ textual descriptions,^40,41^ side effects,^42,43^ drug response,^44^ and drug ontology.^45,46^ We combined these eight modalities of a single drug to create 64 augmented modality pairs for a pair of drugs, effectively expanding the number of training data instances of the original data 64 times (**Fig. 1b**). Moreover, another advantage of our framework is that we don’t require to see every modality for every drug. Traditional approaches that can consider different modalities need to either collect or impute every modality for every drug. In contrast, we circumvent the missing modality challenge by treating a modality pair instead of a drug pair as a data instance, thus allowing us to train the model just using available modality pairs.

**Fig. 1.**
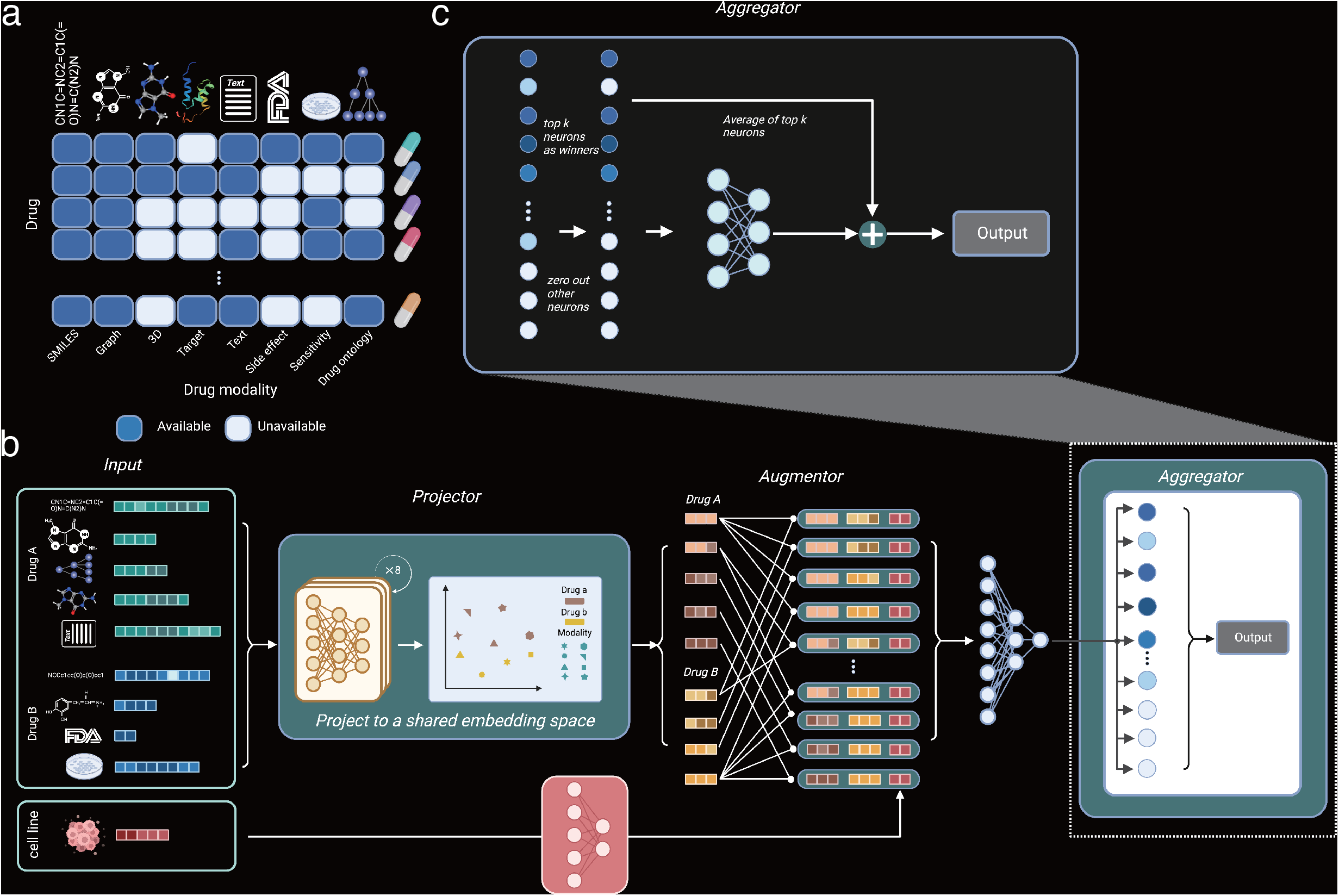
Pisces model overview. **a**, Pisces considers eight different modalities for a single drug, including Simplified Molecular Input Line Entry System (SMILES), molecular graphs, three-dimensional molecular structures, drug targets, textual descriptions, side effects, drug response and drug ontology. Pisces can handle missing modalities by only training on available modality pairs, circumventing the need to impute missing modalities. **b**, Pisces takes a pair of drugs and a cell line as input. Each drug is represented by its available modalities, which could vary between two drugs. Pisces has three components: projector, augmentor and aggregator. The projector trains eight different encoders for eight modalities to project all modalities of all drugs into a shared embedding space. Each point in this space is one modality of one drug, allowing us to expand the original single drug data at most eight times. The augmentor then creates embeddings of a drug pair through the pairwise combination of each drug’s embedding, allowing us to expand the original drug pair data approximately 64 times. **c**, Pisces predicts one score for each of the 64 views, producing 64 predicted scores for a drug pair on a cell line. The aggregator first finds the top *k* largest predictions and zero-out the remaining ones, allowing us to exclude the noise from low-quality augmentations. It then uses a ResNet structure to integrate these *k* largest predictions as the final prediction.

We systematically evaluated Pisces on three drug combination tasks: cell-line-based drug synergy prediction, xenograft-based drug synergy prediction, and drug-drug interaction prediction. On GDSC-combo,^47^ which is a recently-released cell-line-based drug synergy dataset, Pisces achieved at least 21.4%, 23.8%, 10.2% improvements on *F*_1_ score on three different data split settings compared to five comparison approaches.^19–23^ We further demonstrated how Pisces obtained 0.8525 AUROC on three-drug combination synergy prediction when only trained on two-drug data. By interpreting Pisces’s prediction using a genetic interaction network,^48^ we identified a breast cancer (BRCA) drug-sensitive pathway from BRCA cell lines in GDSC-combo. We validated this pathway on an independent BRCA tumor dataset^49^ and found that patients with this pathway have substantially longer survival time (log rank *p*-value < 0.02). Our second validation is a large-scale xenograft dataset.^50^ We found that Pisces again outperforms other approaches on predicting minimum tumor volume changes. Furthermore, we showed that Pisces can accurately predict the tumor volume at an unmeasured time point with a Spearman of 0.47, while existing approaches cannot be applied to such temporal analyses. We visualized that the embeddings learnt by Pisces are ordered in chronological order and matched well with tumor volume, supporting clinical decision making and drug resistance detection. Finally, we applied Pisces to drug-drug interaction prediction, where the goal is to predict whether two drugs will have a certain interaction or adverse drug reaction. On DrugBank^39^ and TwoSIDES,^51^ Pisces outperformed existing approaches when one or both drugs in each test combination were never-before-seen. By examining drug-drug interactions that significantly occurred between drug classes, we constructed a novel DDI network that can be used to infer interactions for new drugs. Collectively, Pisces effectively models drug pairs through augmenting each drug pair to 64 modality pairs, and can be broadly applied to various biological applications that involve a pair of drugs.

## Results

### Overview of Pisces

Pisces takes a triplet of two drugs and a cell line as input and outputs the predicted synergy for this triplet (**Fig. 1b**). Pisces can handle drugs with different available modalities without imputing missing modalities. Pisces is divided into three components: projector, augmentor and aggregator. The projector will enable different modalities, which have unmatched features, to be comparable. Specifically, the projector learns eight neural networks to embed eight different modalities into a shared embedding space (**Supplementary Figure 1**). Each point in the shared embedding space is one augmented view of a particular drug based on a particular modality. For example, drug A with 5 available modalities will be mapped to five different points, each representing one of its modalities. As a result, the projector allows us to expand the original dataset at most eight times. The augmentor combines the augmented views of single drugs into augmented views of drug combinations. In particular, drug A with 5 modalities and drug B with 4 modalities will be combined into 20 augmented views through a pairwise combination. These 20 augmented views will be combined with cell line features to become 20 new data points. Therefore, the augmentor allows us to expand the original dataset at most 64 times.

The aggregator aggregates the predictions from different augmented views into a final prediction (**Fig. 1c**). Pisces leverages the idea of noisy label learning^52^ by only considering top *k* predictions among predictions from all augmented views for a triplet. Only these top predictions will be used by a ResNet^53^ architecture to calculate the final prediction, allowing Pisces to exclude noise from low-quality augmented views. Pisces can be broadly applied to cell line, xenograft and patient data, and other biomedical applications involving a pair of drugs.

We first investigated the agreement among using different drug modalities for drug synergy prediction. We trained eight drug synergy prediction models (**Fig. 2a**), each only using one of the modalities. On one hand, we observed that two modalities often have a Spearman correlation greater than 0.6 (**Fig. 2b, Supplementary Figure 2-4**), indicating a good agreement among them. On the other hand, none of these Spearman correlations is greater than 0.8, even for the one between SMILES and graphs (**Fig. 2c**), which can be converted to each other programmatically.^54^ This demonstrates the complementary information among these modalities, both from different information captured by each modality and from the different neural network architectures used to convert them into machine-readable embeddings. For example, SMILES is embedded using a Transformer, while the molecular graph is embedded using a graph convolutional network (**Supplementary Figure 1**). The agreement and complement of these modalities raises our confidence about the merit of Pisces and motivates us to evaluate it on large-scale drug synergy datasets.

**Fig. 2.**
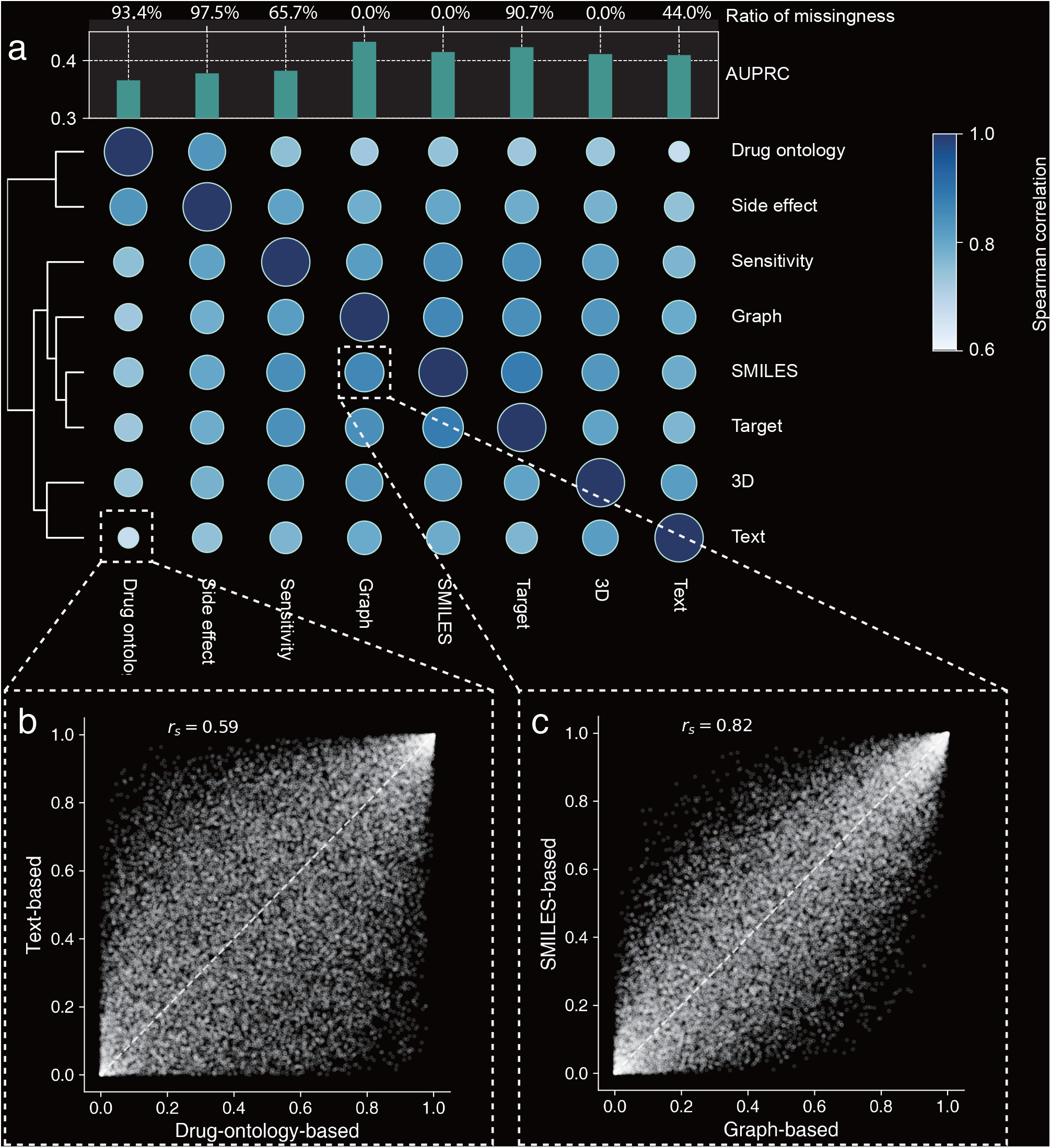
The agreement among predictions of different modalities. **a**, Heatmap showing the agreement between predicted synergy probabilities on GDSC cell lines using the modality in the x-axis and y-axis. Agreement is measured using the Spearman correlation. The same single modality is used for both drugs. Predictions are evaluated using the AUPRC. Ratio of missingness means the percent of drugs that do not have this modality. **b**, Scatter plot comparing the prediction scores of using drug-ontology-based modality and text-based modality. **c**, Scatter plot comparing the prediction scores of using graph-based modality and SMILES-based modality.

### Pisces improve cell-line-based drug synergy prediction

We first sought to evaluate Pisces on cell-line-based drug synergy prediction. We exploited a recently published drug combination dataset from GDSC,^47^ and examined three data split settings: vanilla cross validation, split by drug combination, and split by cell line. On vanilla cross validation, Pisces substantially outperformed five existing approaches on all four metrics (**Fig. 3a-d**). This prominent performance of Pisces indicates the possibility of learning high-quality drug pair embeddings by integrating different modalities. We next examined the split by drug combination setting, where all test drug combinations have never been seen in the training set. Since experimental measurements can only cover a small proportion of all possible drug combinations, this setting can simulate whether our method can predict the responses for new drug combinations. We found that the performance of all methods dropped, indicating that this is a more challenging setting compared to the vanilla cross validation. Nevertheless, Pisces still surpassed the next-best-performing approach by 24% and 26% in terms of the *F*_1_ and KAPPA scores, demonstrating that the embeddings learnt by Pisces on a small collection of training drug pairs can be extended to never-before-seen drug combinations. Finally, we evaluated the split by cell line setting, where all test cell lines have never been seen in the training set. This setting examines whether we can recommend drug combinations to a new patient who has not been tested on any combinations. We found that this setting is the most challenging one among all the three settings according to the overall prediction performance. However, Pisces still obtained 10% improvements in terms of *F*_1_ score against the next-best-performing approach. Collectively, the prominent performance of Pisces on three different settings demonstrates its broad applicability to new drug combinations and new cell lines, confirming the effectiveness of augmenting existing dataset using different drug modalities.

**Fig. 3.**
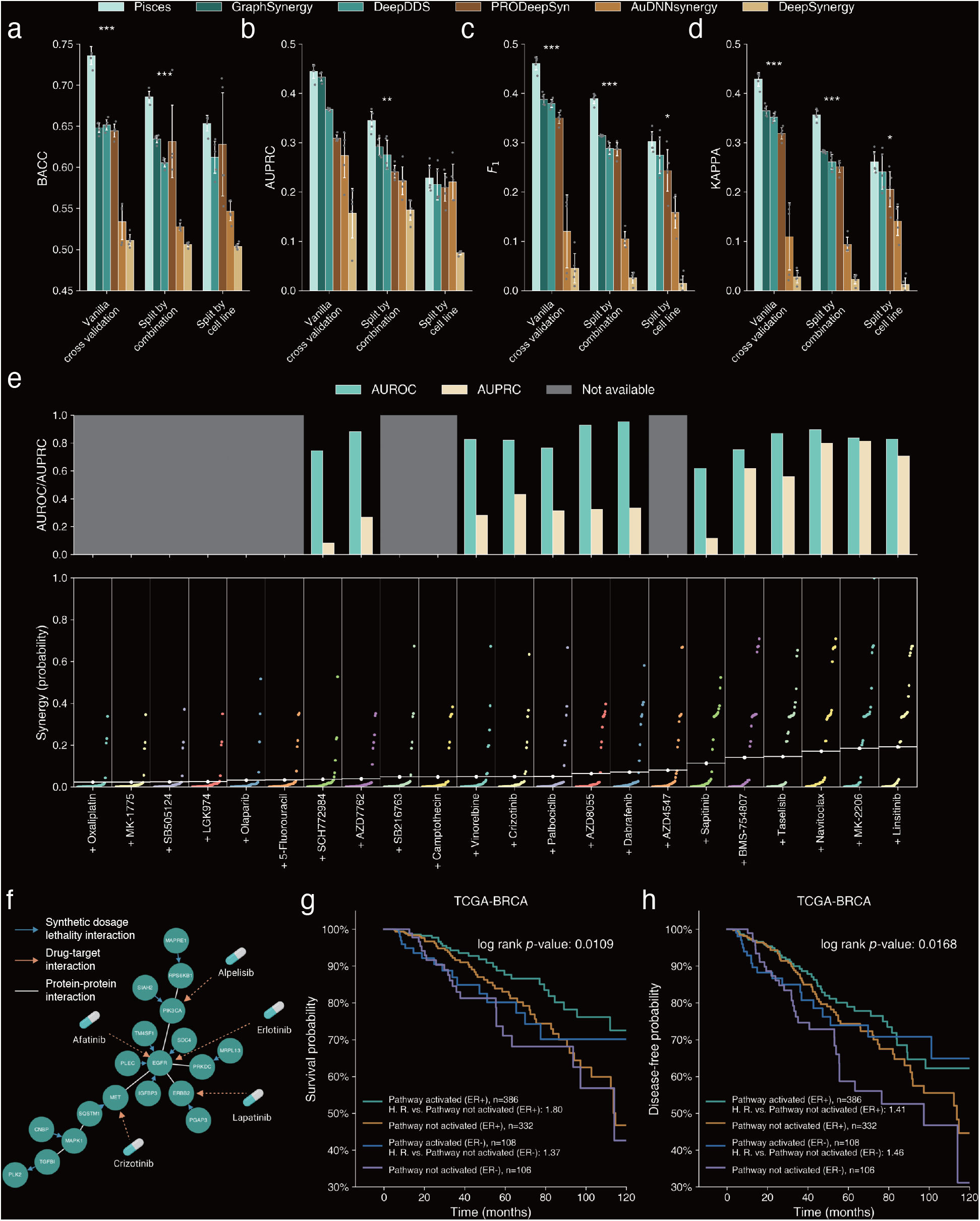
Drug synergy prediction on cell lines. **a-d**, Bar plots comparing the drug synergy prediction on GDSC under three data split settings (x-axis) using BACC (**a**), AUPRC (**b**), F1 (**c**), Cohen’s Kappa (**d**). Split by combination means all test combinations have never been seen in the training data. Split by cell line means all test cell lines have never been seen in the training data. The * indicates that Pisces outperforms the next-best-performing model in the metric, with significance levels of t-test p-value < 5e-2 for *, t-test p-value < 1e-2 for **, and t-test p-value < 1e-3 for ***. All t-tests are one-sided. **e**, Bar plot showing Pisces’s prediction performance on three-drug combination prediction when trained only on two-drug and single-drug data. AUROC and AUPRC are only calculated for three-drug combinations with at least one measured cell line. Afatinib and Trametinib are two of the three drugs and the third drug is shown in x-axis. Each colored point is Pisces’s predicted synergy (y-axis) of this three-drug combination on a cell line. **f**, A BRCA drug-sensitive pathway identified by interpreting Pisces’s drug synergy predictions using a genetic interaction network. **g**,**h**, Survival plots showing the significantly different overall survival (**g**) and disease-free survival (**h**) among four groups of TCGA-BRCA patients. These four groups are classified using the gene expression in the BRCA drug-sensitive pathway and *ER* status.

Existing drug synergy prediction approaches are mainly developed for combinations of two drugs. In practice, combinations of three drugs have been shown to alleviate side effects ^55^ and drug resistance.^56,57^ Pisces can be extended to predict the synergy of three drugs by only training on the combinations of two drugs (see Methods). On the three-drug combination data in GDSC, Pisces obtained an AUROC of 0.82 and an AUPRC of 0.43 on ranking cell lines for a combination (**Fig. 3e**), and 0.88 AUROC and 0.71 AUPRC on ranking combinations for a cell line (**Supplementary Figure 5-6**). Afatinib and Trametinib have been reported to be an effective combination by co-inhibiting of *ErbB* family and *MEK/ERK* pathways.^58^ Pisces predicted that Linsitinib is the most effective third drug when used together with Afatinib and Trametinib on 33 colon adenocarcinoma cell lines (average synergy probability 19.2%). Previous studies demonstrated that Linsitinib could overcome acquired Afatinib resistance by inhibiting *IGF1R* phosphorylation, making the combination of Linsitinib and Afatinib an effective combination therapeutics.^59^ The promising performance of using Pisces to predict the synergy of three-drug combinations indicates that Pisces can be applied to applications that involve more than two drugs.

### Pisces identifies a breast cancer drug-sensitive pathway

To further illustrate the clinical relevance of Pisces, we integrated the predictions of Pisces on 51 BRCA cell lines with a genetic interaction network^48^ to identify a BRCA drug-sensitive pathway (**Fig. 3f**). In particular, since Pisces can be applied to never-before-seen combinations, we first used Pisces to predict the response of 176,344 triplets of cell line and a pair of drugs that have not been experimentally measured in GDSC. Pisces predicted 5.1% of these triplets to be synergistic, demonstrating the rareness of effective drug combinations. We then created 97,479 gene pairs from these synergistic triplets using the following criteria: One gene is over-expressed in the cell line and the other gene is the target of the drug combination. We next filtered these gene pairs using synthetic dosage lethality (SDL) interactions from a genetic interaction network.^48^ The resulting gene pairs are connected as a pathway with 19 genes and 32 edges, including 8 drug targets and 12 SDL interactions. Many genes in this pathway are from *PI3K* (phosphoinositide 3-kinase) and *EGFR/HE*R families. *PI3K* plays a crucial role in *ER*+ (estrogen receptor-positive) tumors that have developed resistance to hormonal therapy.^60,61^ At least 14 drugs in GDSC can inhibit one of these 8 targets (**Supplementary Figure 7**). We hypothesize that this is a breast cancer drug-sensitive pathway since cancer drugs applied to this pathway will inhibit their targets and exploit SDL to effectively kill the cancer cells.

We validated this hypothesis on BRCA tumors in TCGA.^49^ We clustered 977 BRCA tumors into four clusters based on whether this pathway was activated or not and Estrogen receptor (ER) status. We found that *ER*+ tumors that activated this pathway had substantially longer overall survival time (log rank *p*-value < 0.011) and disease-free survival time (log rank *p*-value < 0.017) (**Fig. 3g-h**). To further confirm that the survival prediction ability is not simply from clustering patients to *ER*+ and *ER*-, we clustered 977 BRCA tumors into two groups based on the activation of this pathway. We found that the activation of this pathway still enabled a significantly longer overall survival time (log rank *p*-value < 0.003) and disease-free survival time (log rank *p*-value < 0.008) (**Supplementary Figure 8**), reflecting the possibility to use this pathway as clinical biomarkers for breast cancer prognostic.

### Pisces improves xenograft-based drug synergy prediction

After observing the promising performance of Pisces on predicting drug synergy for cell lines, we next validated it on patient-derived tumor xenografts. We exploited a xenograft dataset that spans 1,238 drug combinations across 277 xenograft models. We first used Pisces to predict the BestResponse, which is defined as the minimum tumor volume change after 10 days, for a pair of drugs on a xenograft model in the cross validation setting. Pisces outperformed other drug synergy prediction approaches using both Spearman correlation and Pearson correlation (**Fig. 4a, Supplementary Figure 9-10**), demonstrating the promising performance of using Pisces to predict tumor growth.

**Fig. 4.**
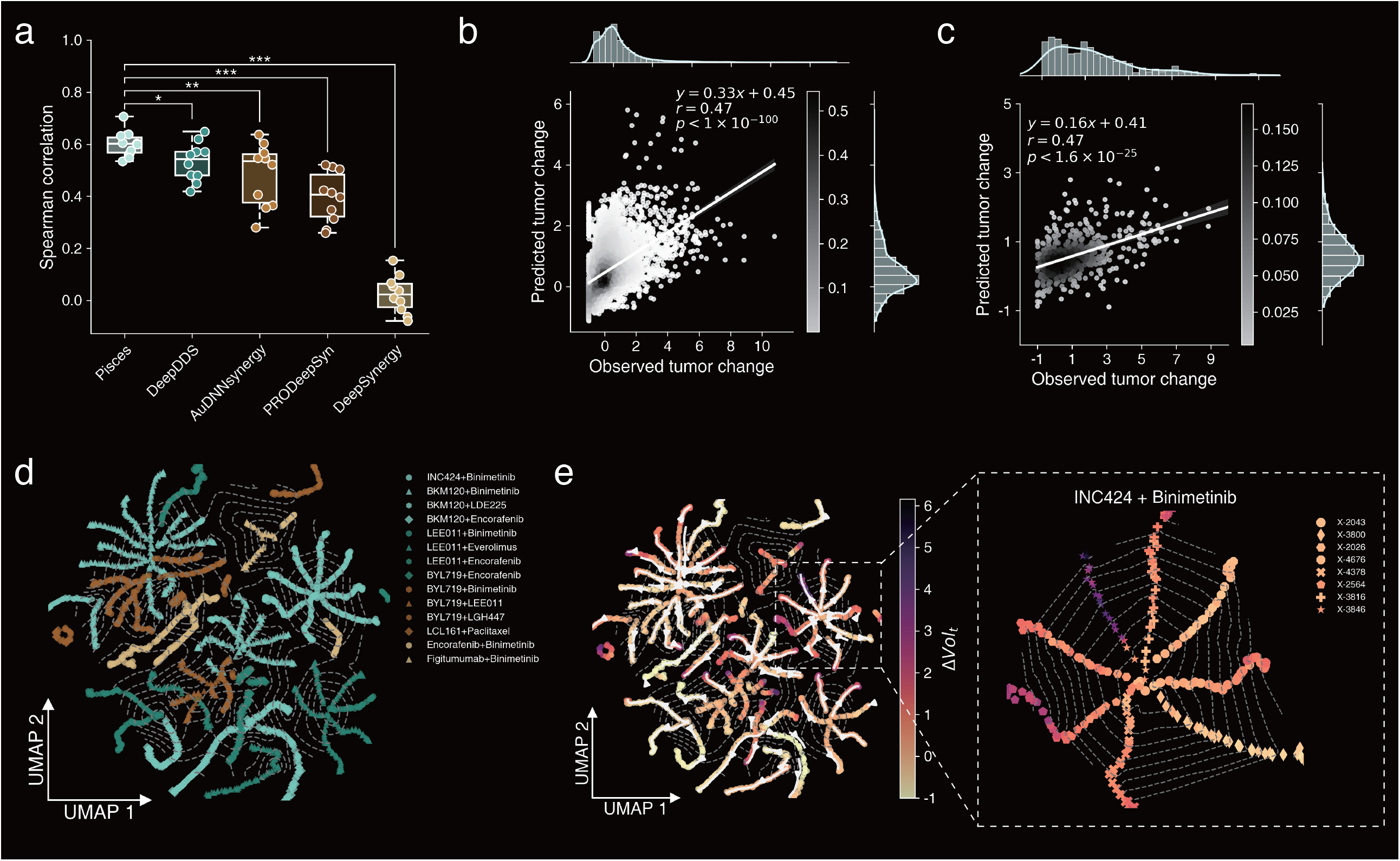
Drug synergy prediction on xenografts. **a**, Box plot comparing the drug synergy prediction on xenografts using Spearman correlation. The * indicates that Pisces outperforms the next-best-performing model in the metric, with significance levels of t-test p-value < 5e-2 for *, t-test p-value < 1e-2 for **, and t-test p-value < 1e-3 for ***. All t-tests are one-sided. **b**, Scatter plots comparing the predicted tumor change and the observed tumor change at holdout time points. Bar plots show the frequency of points in the scatter plot. **c**, Scatter plots comparing the predicted tumor change and the observed tumor change when only the last time point is held out. Bar plots show the frequency of points in the scatter plot. **d**, Each point in the UMAP plot is the embedding of a particular pair of drugs on a particular cell line at a particular time point. Points are colored and marked according to the drug pair. The contours connect tumors that have the same time point. **e**, Each point in the UMAP plot is the embedding of a particular pair of drugs on a particular cell line at a particular time point. Points are colored and marked according to the drug pair. The contours connect tumors that have the same time point. The arrows are from early time points to later time points.

Moreover, predicting tumor growth at a future time point is important for cancer prognosis and survival prediction. Therefore, we extended Pisces to perform such temporal prediction for tumor growth. In particular, we incorporated a time embedding into our framework so that Pisces can now take the time point as an additional input. Pisces can then predict the tumor growth of a drug combination on a xenograft model at any unobserved time point. We found that Pisces obtained a 0.47 Pearson correlation on predicting tumor growth at holdout time points (p-value < 1^*^10^−100^) (**Fig. 4b**). Moreover, we examined an extrapolation setting, where only the last time point of each xenograft model is held out, and observed that Pisces also accurately predicted the tumor growth (Pearson *r* = 0.47, p-value < 1.6^*^10^−25^) (**Fig. 4c**), demonstrating its applicability to perform temporal prediction on tumor growth.

After observing Pisces’s prominent performance on temporal tumor growth prediction, we are motivated to examine the tumor growing trajectories using embeddings generated by Pisces. To this end, we visualize the embedding of test triplets (i.e., a drug combination on a xenograft model) in the two-dimensional space. We first found that these triplets are clustered by xenograft models and branches in each cluster represent different drug combinations (**Fig. 4d, Supplementary Figure 11**). By coloring each triplet using time points, we find that each branch grows from an earlier time point to a later time point, reassuring Pisces’s ability to capture the temporal tumor growth. By coloring each triplet using tumor volume change and adding contours to characterize time points, this embedding space can illustrate both tumor growth and time point, supporting the identification of drug resistance (**Fig. 4e, Supplementary Figure 12**). For example, INC424 and binimetinib showed diverse drug responses among different xenograft models, where the tumor grows most aggressively on X-3846 and X-2564, suggesting drug resistance. Collectively, the promising performance of Pisces on xenograft indicates its applicability to the *in vivo* setting, as well as unique tumor temporal analysis that can hardly be achieved by existing drug combination prediction methods.

### Pisces accurately predicts drug-drug interactions

The promising results of Pisces on drug synergy prediction further motivates us to apply it to other tasks that involve a pair of drugs. To this end, we investigated whether Pisces can be used to predict drug-drug interactions (DDI), where the goal is to classify a pair of drugs into hundreds of predefined interaction types. We evaluated our method on two large-scale DDI datasets: DrugBank, which contains 191,402 drug pairs over 86 interactions, and TwoSIDES, which contains 63,472 drug pairs over 963 interactions. On DrugBank, we found that Pisces substantially outperformed six existing approaches under three different settings, including vanilla cross validation, one new drug in each test pair and two new drugs in each test pair (**Fig. 5a-c**). The improvement against competing methods is larger under the more challenging settings of one new drug in each test pair and two new drugs in each test pair, indicating the strong generalizability of Pisces through integrating different modalities. On a larger dataset TwoSIDES, Pisces outperformed five comparison approaches and achieved a comparable performance with R^2^-DDI (**Fig. 5d-f**), again highlighting its broad applicability to drug-drug interaction prediction.

**Fig. 5.**
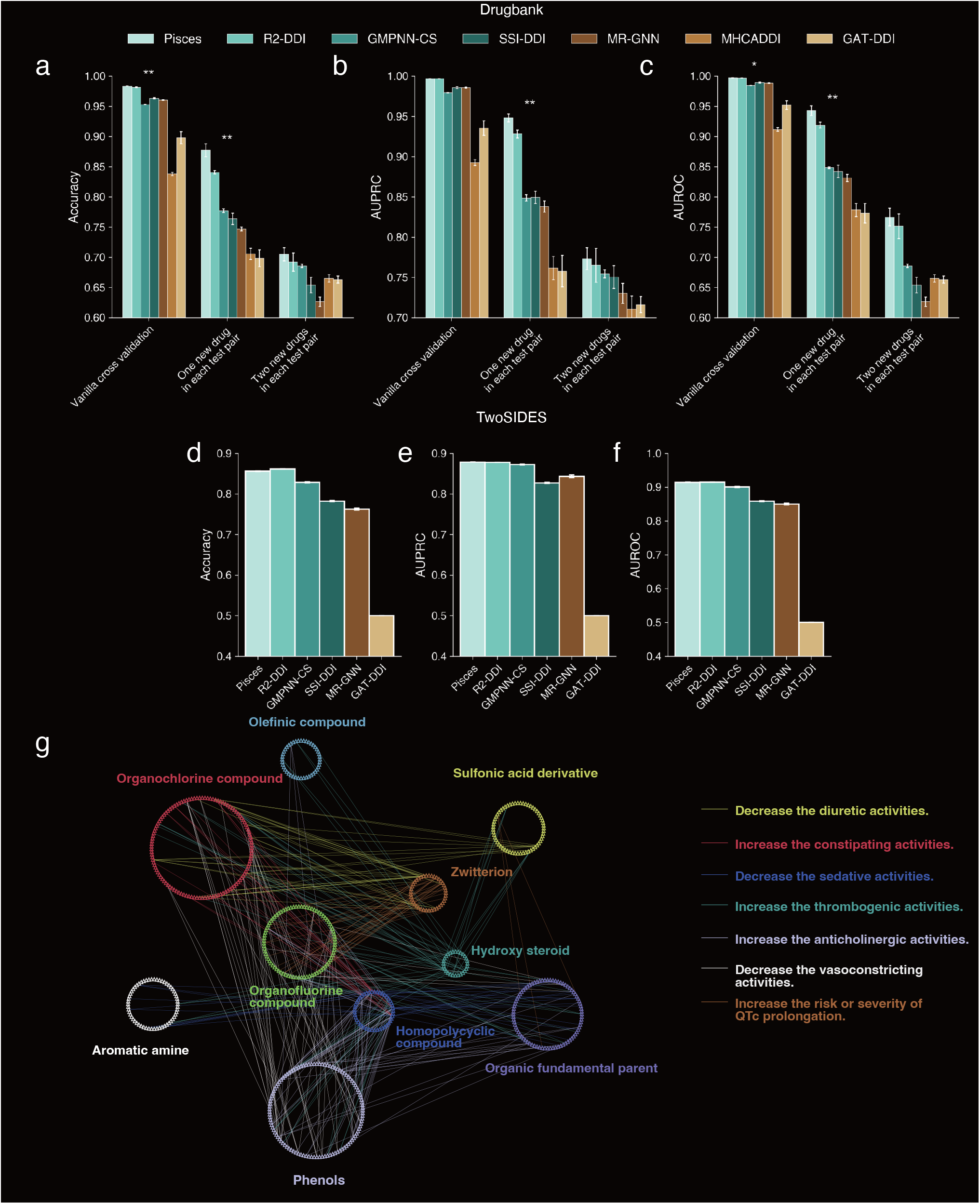
Drug-drug interaction predictions on DrugBank and TwoSIDES. **a-c**, Bar plots comparing the prediction performance of drug-drug interactions on DrugBank under three cross validation settings using Accuracy (**a**), AUROC (**b**), and AUPRC (**c**). One new drug in each test pair means that one and only one of the two drugs in a test drug pair has never been seen in the training data. Two new drugs in each test pair means that both of the two drugs in a test drug pair have never been seen in the training data. **d-f**, Bar plots comparing the prediction performance of drug-drug interactions on TwoSIDES using Accuracy (**d**), AUROC (**e**), and AUPRC (**f**). **g**, Visualization of significantly occurred interaction types between two drug classes determined by Fisher’s exact test (p-value < 0.05). Nodes are colored by drug classes in the drug ontology. Edges are colored by interaction types.

Finally, we examined DDIs that are not collected in DrugBank, but are predicted with high confidence by Pisces. We first group drugs based on their classes in the drug ontology. We then used Fisher’s exact test to find interaction types that significantly occurred between two drug classes (**Supplementary Table 1**). We form these significant interactions as a DDI network, where nodes are drugs and edges are colored by the most likely interaction between two drugs (**Fig. 5g**). This novel DDI network offers a global view of DDI and can extend our analysis to never-before-seen drugs according to its class in the drug ontology.

## Discussion

We have proposed Pisces, a novel machine learning approach for drug synergy prediction. Pisces can augment existing sparse drug combination dataset at most 64 times using 8 different modalities of each drug. We have shown the prominent performance of Pisces on cell-line-based drug synergy prediction, xenograft-based drug synergy prediction, and drug-drug interaction prediction tasks. We also demonstrated the interpretability and clinical relevance of Pisces by identifying a breast cancer drug-sensitive pathway, and verified using survival data from TCGA. Collectively, Pisces will facilitate the future discovery of drug combinations and drug-drug interactions, and can be applied to a wide range of biological applications that involve multiple drugs.

Pisces has at least two limitations that we would like to address in future works. First, the large number of modality pairs increase the predictive performance, but also makes it hard to understand what modalities are most important for a certain prediction. We plan to exploit local interpretation methods to interpret the prediction results.^62,63^ Second, while we have considered different modalities as features, we didn’t consider different types of labels jointly. Labels for drug pairs include synergy effects over cell lines and xenografts, side effects, and interactions. Since our framework can handle missing modality, it can also be extended to handle missing labels. We plan to incorporate multi-task learning^64^ into our framework so that Pisces can be simultaneously optimized for different biological tasks that involve a pair of drugs.

There are a few existing cross-modal drug representation learning approaches,^38,65–68^ such as MolCLR^66^ and the geometry enhanced GEM^65^. There are at least two key differences contrasting Pisces with these works. First, these existing cross-modal approaches mainly study two drug modalities, such as SMILES and molecular graphs. In contrast, Pisces integrates eight different modalities, including structure features, mode of action features, phenotypes, and pharmacodynamic features, allowing Pisces to learn much more comprehensive embeddings for drugs. Second, these approaches only model a single drug, while Pisces is designed for a pair of drugs. Our pairwise combination of single drug augmentation is a novel approach to augment a pair of drugs, effectively augmenting existing sparse datasets up to 64 times. Compared to existing drug combination prediction approaches,^19–23^ Pisces also has at least three key differences. First, Pisces exploits novel machine learning techniques, including data augmentation, contrastive learning and noisy label learning, which alleviate the sparsity and noise in drug combination dataset. Second, existing drug synergy approaches were designed specifically for synergy prediction, while Pisces is a general framework for tasks that study a pair of drugs. Third, Pisces can be applied to combinations with more than two drugs, while existing approaches can only be applied to two-drug combinations, demonstrating the broad applicability of Pisces.

## Methods

### Problem setting

Let Ɗ be the set of drugs and *C* be the additional features. The input drug combination dataset was defined as multiple triplets 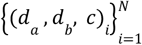, where *d*_*a*_ ∈ Ɗ and *d*_*a*_ ∈ Ɗ is a pair of drugs. We used *c* ∈ *𝒞* to represent the additional features of this drug pair. *𝒞* can be the cell lines and the xenograft models in drug synergy prediction tasks or the indication of interaction types in DDI prediction. Both tasks can be modeled as the binary prediction task:

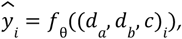

where *f* _θ_ : Ɗ × Ɗ × *𝒞* is a learned mapping function with θ as parameters. The output 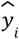 denotes the predicted property of the input triplet (*d*_*a*_, *d*_*b*_, *c*)_*i*_ being positive, which can be the probability of being synergistic in cell-line-based drug synergy prediction, the tumor volume change in xenograft-based drug synergy prediction and the probability of the drug pair exhibiting a specific interaction type in drug-drug interaction prediction.

We considered 8 different feature modalities for each drug: the SMILES, molecular graphs, drug target genes, drug three-dimensional structures, textual descriptions, drug ontology views, drug side effects and drug sensitivity, which we used

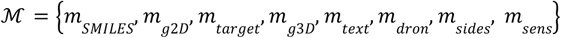

to denote them respectively. We denote the input sample with a pair of modalities as *x*[*m*_*a*_, *m*_*b*_] = (*d*_*a*_[*m*_*a*_], *d*_*a*_[*m*_*a*_], *c*), where *m*_*a*_, *m*_*b*_ ∈ ℳ. Considering |ℳ| different modalities, each drug combination produces a set of |ℳ| × |ℳ| modality pairs {*x*[*m*_*a*_, *m*_*b*_]|∀*m*_*a*_, *m*_*b*_ ∈ ℳ}. By treating each modality pair as an augmented drug combination view, we now expand the original data by |ℳ| × |ℳ| times.

### Projector with modality-specific drug encoders

The eight modalities have different data structures. In order to integrate them together, we designed a projector module that transformed these modalities into a shared space, e.g. dense embeddings with the same length. The diverse data structures of different modalities motivate us to learn a set of embedding functions {*f*_θ[*m*]_ |*m* ∈ ℳ} separately with parameters θ[*m*] for |ℳ| modalities.

### 1. SMILES modality

Each drug SMILES string can be represented as *m*_*SMILES*_ = ⟨*s*_1_, …, *s*_*l*_ ⟩ where *s*_*i*_ denotes the *i*-th token of the *l* length string. Since Transformer has emerged as an efficient model to embed a wide range of sequence features, such as natural languages,^31,32^ protein sequences^69,70^ and molecular features,^71^ we adopted the encoder part of Transformer as 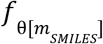. We appended the special token [*CLS*] to each SMILES string: 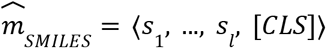 and get the contextualized embedding at the position corresponding to the special token as the embeddings 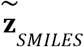 for this SMILES string. We finally used one transformation *MLP* layer to get the final embedding: 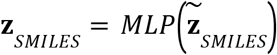, where 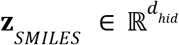.

### 2. 2D graphs

Each molecule can be represented as a 2D molecular graph *m*_*g*2*D*_ = (*V, E*), where 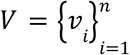 are *n* atoms in this molecule and *E* are the set of bonds between atoms.

We used DeeperGCN^72^ as the graph encoder 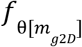 to calculate embeddings for each atom: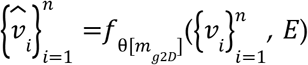. Finally, we used mean pooling and max pooling to get the graph embedding: 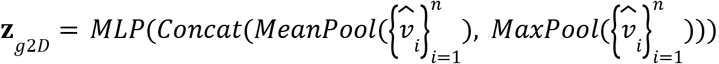 where 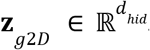

### 3. Drug target

We denote the target gene set of each drug as 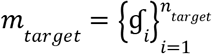. To utilize the physical and functional interactions between genes, we find the *k*_ɠ_ nearest neighbors for each ɠ ∈ *m*_*target*_ in the protein-protein interaction networks (PPI), defined as 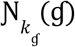 and finally compute the expanded gene set 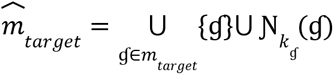. For each gene 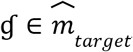, we used one learnable embedding as its representation: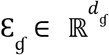. We finally compute the drug target embedding using an *MPL* layer 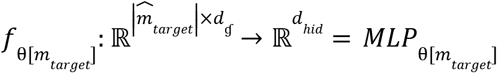 as:

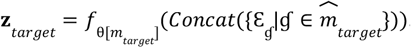

### 4. Drug 3D geometric views

We also sought to learn the drug representations from 3D geometric views. By projecting the drug 2D graph modalities towards the 3D geometric views, we are able to enhance each drug with 3D information. Let Ф_2*D*_ (*V, E*) be the 2D graph encoder and Ф_3*D*_ (*V, O*) be the 3D geometric encoder, where *O* denotes the 3D coordinates of each atom. The 2D graph encoder is aligned with the 3D encoder using a contrastive learning loss:

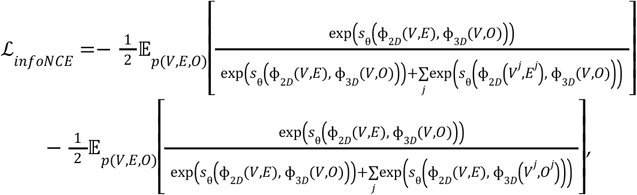

where *s*_θ_ is the scoring function for computing similarities between 2D and 3D views and *j* corresponds to the randomly sampled complementary views. Using the fixed 2D graph encoder aligned with 3D view, we could get the drug embedding augmented by both views. We use our encoder 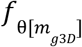 to calculate embeddings:

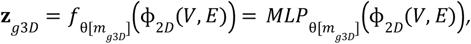

where 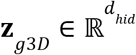 represents the output embedding with 3D geometric views. In our implementation, we utilized the pretrained GraphMVP^38^ as the Ф_2*D*_ (*V, E*) encoder.

### 5. Drug textual descriptions

For each drug, we used their textual descriptions from the ChEBI dataset. For drugs that are not collected in ChEBI,^41^ we used MolT5,^40^ a translation framework from molecules to natural languages, to generate their textual descriptions. We denote the each textual description as a token sequence 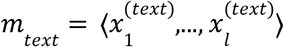. To learn a dense representations from textual descriptions, we used a parameter-fixed biomedical domain specific pretrained language model PubMedBERT^73^ together with an *MPL* mapping function as the encoder to embed textual descriptions for each drug 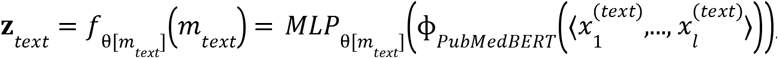, where 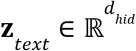 represents the output text modality embeddings.

### 6. Drug ontology structures

Drugs can be queried as a node in the drug ontology, which is a structured controlled vocabulary to classify drugs. We followed Mashup^74^ to get node embeddings. We constructed the transition matrix **𝒯** from the ontology structure using the Random Walk with Restart (RWR):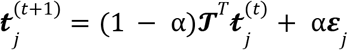 where 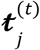 denotes the probability vector of transiting from node *j* to all nodes. α is the restart probability and ***ε***_*j*_ is the basis vector. Then we got node embeddings by singular value decomposition: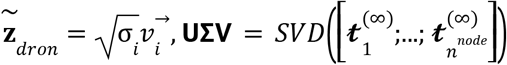. Let’s use 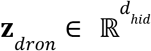 denote the ontology embeddings. Finally, we got **Z**_*dron*_ using our drug ontology encoder:

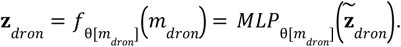

### 7. Drug side effects

We collected the drug side effects from the SIDER 4.1 dataset^42,43^ which includes 1,428 drugs and 27 side effect types. For each drug, we used the one hot embedding 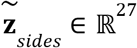 to represent their side effects, where 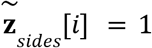 means this drug has the *i*-th side effect. We transformed the side effects into dense embeddings 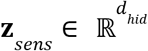 using: 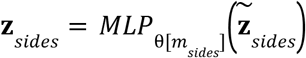.

### 8. Drug sensitivity

The drug sensitivity data was the drug response measured across NCI 60 cell lines.^44^ Each drug response can be represented as 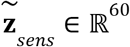 where each element represents the response of a single cell line. We transformed 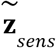 to get the drug sensitivity representations 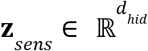 using one *MLP* layer. The transformation function can be denoted as 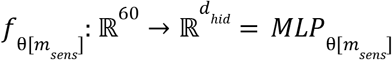.

This projector module enables us to embed each drug with |ℳ| modalities into |ℳ| dense embeddings. Next, we sought to learn representations for a pair of drugs.

### Augmentor for creating augmented drug combinations

The |ℳ| modalities enable us to obtain |ℳ| × |ℳ| views for a pair of drugs, resulting in augmented drug combinations. For the drug combination triplet (*d*_*a*_, *d*_*b*_, *c*_*i*_), if we use the embedding **Z**_*m*_ from a single modality *m* ∈ ℳ to represent one drug, the embedding of additional feature 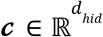 to represent the additional features *c*, we were able to combine them into |ℳ| × |ℳ| triplets: 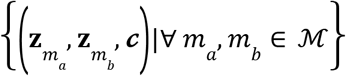. The augmentor module applied one transformation function to generate augmented drug combination views:

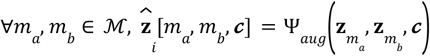

Note that with these new data instances, our training dataset is |ℳ| × |ℳ| times larger. Passing all augmented drug combination views through a shared classifier, we were able to collect |ℳ| × |ℳ| predictions for each triplet. Let’s denote the classifier as 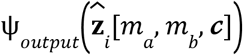, and the collected predictions for triplet (*d*_*a*_, *d*_*b*_, *c*)_*i*_ is 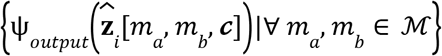.

### Aggregator module for effective aggregations of all predictions

For each drug triplet, with the projector and augmentor modules, we now collected |ℳ| × |ℳ| predictions from augmented drug combination views. Finally, we aggregated the output predictions together. Inspired by the noisy label learning, we sought to consider the top predictions as highly confident predictions and treat others as predictions from low-quality augmented views. We set *k* data instances with larger prediction values to be kept while other data instances to be ignored. One *NLL*() layer was added to perform this operation. This layer basically keeps the top *k* neurons with higher prediction values while setting the output of other neurons to be zero. Then we applied a linear affine matrix ***ω***_*NLL*_ together with a shortcut to aggregate the top predictions. Taken together, the output of this aggregator module can be calculated using the following equation:

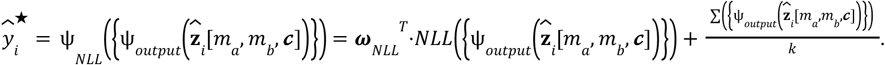

### Training objective of Pisces

The training objective of Pisces consisted of two parts: the supervised training objective and the consistency objective. The supervised training objective aims to minimize the loss between the ground truth label and the predictions either from each augmented data instance or from the aggregated predictions. We denote the supervised training objective as:

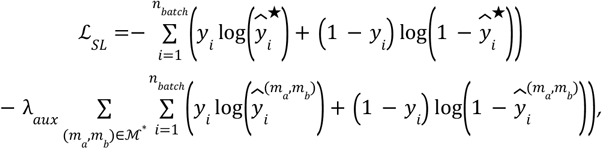

where *y*_*i*_ denotes the ground truth label of the *i*-th sample and λ_*aux*_ is a hyperparameter. We also used 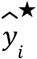 to denote the aggregated output and used 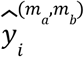 to denote the output when drug *a* and drug *b* use *m*_*a*_ and *m*_*b*_ modalities respectively. In the second term, we used a sampling strategy to reduce the computational costs. We used 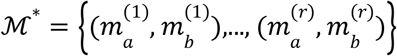 to denote a set of modality pairs uniformly sampled from all combination set ℳ × ℳ. In our experiments, we set *r* = 2.

Since the augmented data instances from the same drug triplet are not statistically independent, we introduced a consistency objective to encourage these instances to correlate with each other. We considered these correlations in terms of both the embedding space and the output space. A contrastive learning loss was introduced to improve the consistency among augmented views in terms of the embedding space. We adopted the InfoNCE loss,^75^ which has been broadly applied in diverse machine learning areas,^76–79^ to improve correlations among augmented views within one batch:

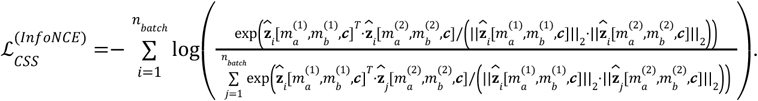

In addition to the contrastive learning loss at the embedding space level, we added another loss term to improve consistency in the output. We applied the mean squared error (MSE) loss to let the aggregated prediction 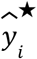 instance: to be correlated with prediction from each augmented data

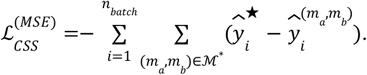

Inspired by noisy label learning,^52,80,81^ the key intuition of this consistency loss is to exploit high-quality refined prediction 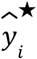 as a soft label to correct the rest of noisy predictions using each modality pair 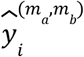. If 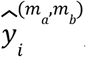, the noisy predictions do not agree with the refined soft label, we encourage the model to adjust them to be close to the refined soft labels. This consistency loss benefits the model to have predictions with high fidelity. Finally, we combined the supervised training loss and consistency loss together as the training objective:

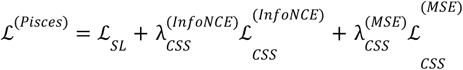

### Collect multi-modalities of drugs

We assume that the drug combination dataset provides the SMILES modalities for all drugs. Graph modalities can be converted from the SMILES modalities using the RDKit python package.^54^ We collected the drug target modalities from both the Genomics of Drug Sensitivity in Cancer (GDSC)^14^ and the DrugBank dataset.^39^ The GDSC dataset contains 621 drugs with target genes and the DrugBank dataset has 14,398 drugs. We aligned them with existing drugs using their names and synonyms to obtain drug targets. To collect the 3D geometric views, we used a 2D graph encoder from GraphMVP, pre-trained with 3D chemical structures using contrastive learning loss, to project our molecules into the representation space augmented with 3D information. The textual descriptions are collected with the MolT5 code base, which both curated textual descriptions for molecules in ChEBI dataset and provides the machine translation tool to generate texts for molecules outside the ChEBI dataset. The drug ontology dataset, with 8,248 drugs in total, was obtained from the National Center for Biomedical Ontology.^82^ We used the version released by October 22, 2022. The drug side effects were collected from SIDER v4.1 dataset^42,43^ and we finally obtained 1,428 drugs and 27 side effect types. The drug sensitivity data, which measure 20,861 drugs across NCI 60 cell lines^44^ in terms of average z score was obtained from the CellMiner website (https://discover.nci.nih.gov/cellminer/loadDownload.do).

### GDSC-combo dataset processing

The GDSC-combo dataset was a recently published Genomics of Drug Sensitivity in Cancer dataset.^47^ The original dataset presents the samples formulated by (Drug A, Drug B, cell line, synergy or not) tuples, and we view each triplet (drug A, drug B, cell line) as synergy if there exists one, otherwise no synergy. We consider the drug combination to be synergistic as long as it is indicated as synergistic under one circumstance. We chose samples that have a combination of 2 drugs and finally obtained 102,893 samples, including 63 drugs and 125 cell lines. We evaluated three settings: vanilla cross validation, split by drug combination and split by cell line settings. In the vanilla cross validation setting, we randomly split the dataset by 60% for training, 20% for validation and 20% for testing. In the split by drug combination cross validation setting, we split the dataset according to drug pairs. We used the splitting ratio of 60%, 20% and 20% for training, validation and testing to split the drug combinations, and distributed samples to different sets based on the drug pairs. We used the same way to split the 125 cell lines features in the split by cell line cross validation setting and collected samples into train, validation and test set.

### Comparison approaches for drug synergy prediction on cell lines

We compared Pisces to five existing drug synergistic prediction approaches. **PRODeepSyn**^23^ takes molecular fingerprints and descriptors for drugs as the inputs. The molecular fingerprint is a 256-dimensional binary vector for each drug, representing the existence of a set of predefined substructures.^83^ Then drug descriptor is a 200-dimensional real vector used representing molecules’ physical or chemical properties of interest, such as lipophilicity or molecular refractivity.^84^ Both the fingerprints and the descriptors are obtained from RDKit.^54^ Cell lines are embedded using Graph Convolutional Networks^85^ by integrating the protein-protein interaction (PPI) network with the gene expression vector. Finally, PRODeepSyn then used an MLP to predict drug synergy. **AuDNNsynergy**^22^ utilizes the same features as those in PRODeepSyn. Different from PRODeepSyn, it trains three autoencoders to predict drug synergy. **DeepSynergy**^20^ takes molecular fingerprints, descriptors and drug-target interactions for drugs and Transcripts Per Million (TPM) for cell lines as the inputs. All these features are then fed into an MLP to predict drug synergy. **GraphSynergy**^21^ utilizes Graph Convolutional Networks to extract drug and cell line features from the PPI network. These features are then fed into an MLP to predict drug synergy. **DeepDDS**^19^ takes molecule graphs for each drug and TPM for cell lines as the inputs. It then uses an MLP to predict drug synergy. Notably, these comparison approaches often rely on very different features that might not be available in any dataset. The original implementations of these methods often use hard-coded or pre-processed features that cannot be generalized to GDSC-combo. The details of pro-processing are also not comprehensively revealed and make it hard to reproduce their results. Therefore, we have re-implemented many of them and use the same feature preprocessing to increase the usability for fair comparison. We have made our implementations of all five comparison approaches available for future studies.

### Implementation details for cell-line based drug synergy prediction task

In the drug synergy prediction, the input of our model is a triplet, including a pair of drugs and the cell line feature, and the output is the probability of the drug pairs being synergistic on this cell line. Following the previous works,^19–23^ we used a threshold of 0.5 to determine the class label for the input triplet *x* = (*d*_*a*_, *d*_*b*_, *c*_*i*_). The additional features are gene expression values of one cell line in terms of TPM (transcripts per million), which is a fixed-size vector 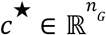 representing gene expression levels across *n*_*G*_ genes. To model the cell line features, we first determined an overexpressed gene set from *c*^★^. We denote this gene set as 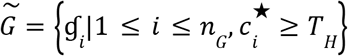 where *T*_*H*_ is a threshold to determine whether a gene is overexpressed in this cell line. In practice, we set *T*_*H*_ = 400. Similar to the processing of the drug target modality, we also added one-hop information on the PPI network: 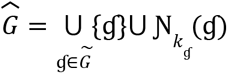 Then the cell line features, which is the context features embeddings, were calculated by aggregating embeddings of all genes in 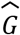 together: 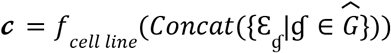 where Ɛ_ɠ_ is learnable embedding for each gene and shared with the drug target modality processing. In the augmentor, we first concatenated the drug embeddings and context features together: 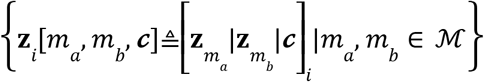. We used one layer MLP as the augmentor transformation function :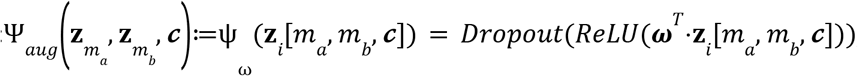, where 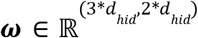. Here 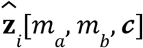 refers to the embeddings for each augmented drug combination. Finally, we applied a classifier to get the predicted probability for each data instance. We denote this classifier as: 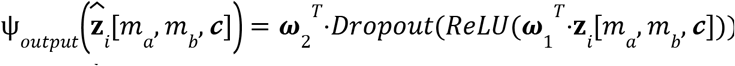, where 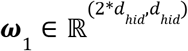 and 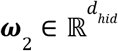.

In the drug synergy prediction task, we set the number of layers of both the DeeperGCN and Transformer to 6 and the hidden dimension size to 384 and 512 respectively. We set the transformed hidden dimension *d*_*hid*_ for all the modalities to 512. The architecture of transformation layers for all modalities {*MPL*_θ[*m*]_ |*m* ∈ ℳ} is one linear layer mapping function with different input dimension sizes. We trained the model using the Adam optimizer with a batch size of 128 for 100,000 steps. We adopted a polynomial decay scheduler with 4,000 warm-up steps. We performed a grid search for the learning rate within [1 × 10^−5^, 5 × 10^−5^, 8 × 10^−5^, 1 × 10^−4^, 5 × 10^−4^]. 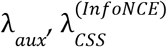 and 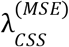 were set to 0.01. We chose top *k* = 8 predictions in the aggregator. To evaluate all comparison approaches and our method on this highly imbalanced dataset, we used four metrics: balanced accuracy (BACC), that is the average of sensitivity (true positive rate) and specificity (true negative rate), area under the precision-recall curve (AUPRC), *F*_1_ score and Cohen’s Kappa statistic. All metrics are higher the better.

In **Fig. 2**, we investigated the agreement between different modalities on the GDSC-Combo dataset. Here we trained eight models based on Pisces’s architecture and each of them only uses one type of modality. For missing modalities, we assigned learnable embeddings and optimized them in the training procedure. In **Fig. 3e**, we also extended Pisces that is trained on two-drug combination to three-drug combination prediction. We first separated each three-drug combination into three two-drug combinations, and got a prediction for each of them. Finally, we used the average predictions to be the synergy probability of these three drug combinations. In **Supplementary Fig. 13**, we investigated how the performance varied if we chose different top *k* in the aggregator. We evaluated the vanilla cross validation setting. First, we observed that balanced accuracy (BACC) was sensitive to the value of *k*. We observed that BACC did not change too much when *k* is smaller than 32, indicating our approach was robust to the selection of *k*. Notably, the performance deteriorated a lot in terms of BACC when we aggregated all 64 predictions to calculate the output, demonstrating that our strategy could exclude noisy predictions.

### Identification of a breast cancer drug-sensitive pathway

The GDSC-Combo dataset has 102,893 triplets (a pair of drugs and a cell line) which encompasses 63 drugs and 125 cell lines. These 125 cell lines include breast, colorectal and pancreas cancer cell lines. Each triplet was represented as (*d*_*a*_, *d*_*b*_, *c*)_*i*_. Here *d*_*a*_ is the anchor compound and *d* is the library compound. Notably, (*d*_*a*_, *d*_*b*_, *c*)_*i*_ and (*d*_*b*_, *d*_*a*_, *c*)_*i*_ represent two triplets with the same drug pair and cell line, but the order of anchor and library compounds was swapped. Considering all possible combinations of these 63 drugs and 125 cell lines, there were 488,250 possible triplets. For each triplet (*d*_*a*_, *d*_*b*_, *c*), if neither (*d*_*a*_, *d*_*b*_, *c*)_*i*_ or (*d, d*_*a*_, *c*)_*i*_ appeared in the collected GDSC-Combo dataset, we collected it as a novel triplet. When collecting novel triplets, if (*d*_*b*_, *d*_*a*_, *c*)_*i*_ had already been collected, (*d*_*a*_, *d*_*b*_, *c*)_*i*_ would not be collected. We finally collected 176,344 novel triplets. We applied Pisces to these novel triplets and filtered out triplets with predicted probabilities less than 0.9. For each triplet, we collected the drug target genes 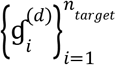 and overexpressed genes 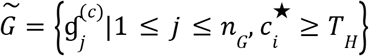 in the cancer cell lines. Therefore, we constructed multiple gene pairs, each pair being represented as 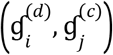 and obtained 97,479 gene pairs in total. For each pair, we treated 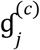 as the overactive genes and the drug target 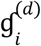 as the essential genes to the cancer cells. This logic matched the genetic interactions in synthetic dosage lethality (SDL) networks.^48^ Therefore, we used a published SDL networks database to filter out gene pairs not in the SDL networks. To focus on studying breast cancer, we selected gene pairs collected from triplets on 51 breast cancer cell lines. We finally collected 12 SDL gene pairs with 19 genes as a breast cancer drug-sensitive pathway.

We verified this breast cancer drug-sensitive pathway on breast cancer patients from the TCGA dataset. We collected the gene mutation data and gene expression data. If the expression level of one gene from a patient was higher than 70% of all patients, we determined this gene as a highly-expressed gene. If one gene was mutated or its expression level was lower than 70% patients, we determined it as inhibited genes. We determined the pathway to be activated as long as there existed one gene pair of the 12 SDL pairs, from which 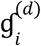 was inhibited and 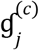 was highly-expressed. We also noticed that the breast cancer patients can be classified into ER+ breast cancer patients and ER-breast cancer patients. Therefore, based on the status of ER and the activation of our identified pathway, we now clustered these breast cancer patients into four groups: pathway activated (ER+), pathway not activated (ER+), pathway activated (ER-), and pathway not activated (ER-) patients. We therefore perform survival analysis using the lifelines python package.^86^

### Xenograft dataset processing

The xenograft dataset refers to the drug response data for the patient tumor derived xenograft models. Hui Gao et al. released the xenograft data for both single drugs and drug combinations that provided tumor volume changes at different time points after transplantation.^50^ We denote the tumor changes at time *t* as 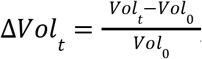, where *Vol*_*t*_ represents the tumor sizes at time *t* and *Vol*_0_ represents the tumor size when *t* = 0. We collected the response of 26 drugs across 191 mice with gene expression features, and 13 drugs of them can find target genes in our collected DrugBank and GDSC drug target dataset. In the raw dataset, the gene expression features were normalized into the FPKM unit. We then converted the FPKM unit into TPM unit:

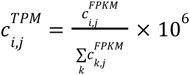

We evaluated 3 settings on the xenograft dataset: (1) We used our model to predict the BestResponse for one drug combination across all time points. The BestResponse means the minimum value for all tumor changes when 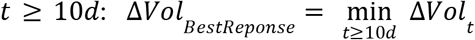. In BestResponse prediction, we finally collected BesResponse of 1,769 single drugs and 476 drug combinations. We used all single drug samples and 60% of the drug combination samples as the training set, and used 20% drug combination samples as validation and test set respectively. For each single drug sample, we replicated them into a drug pair as the input. (2) We evaluated our model’s ability to predict the volume changes at each time point. We also used all the single drug samples as part of the training set and split the drug combination samples into the training, validation and test set. Notably, the drug combinations in all the test samples are never seen in the training set. (3) We applied our model to the extrapolation setting where we only predict the last time point. We evaluated all the three settings using 10 fold cross validation. We also collected 22,156 single drug responses and 9,035 drug combination responses across all time points.

### Implementation details for drug response prediction on xenograft models

The drug response prediction task aims to predict the volume changes for a xenograft model when exposed to a specific drug combination. Since the tumor volume changes represent continuous values, we opted to modify our supervised training objective by replacing the cross entropy loss with the mean squared error loss:

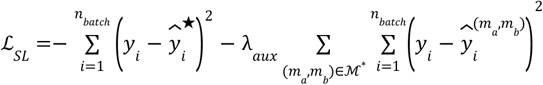

In BestResponse prediction, the input of our model is a triplet (drug A, drug B, model c), where we used the gene expression features ***c***^*exp*^ to represent the model c. We employed the same approach utilized in the drug synergy prediction task to calculate the gene expression embeddings. This gene expression feature embeddings ***c***^*exp*^ were used as the model embeddings.

When predicting drug response at time point *t*, the input of our model can be represented as (*d*_*a*_, *d*_*b*_, *c, t*). Here the time *t* is an integer representing the number of days post to the transplantation. Inspired by the position embedding in Transformer,^87^ we used the following equation to calculate the time embedding:

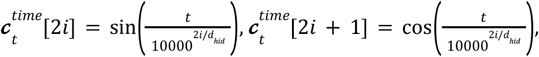

where 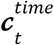 is a fixed length vector with the dimension size of *d*_*hid*_ = 512. In both the drug response prediction at time point *t* and the extrapolation tasks, we concatenated the gene expression embeddings and the time embeddings together as the additional feature embeddings: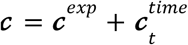.

We used the same hyperparameters of model architectures as the model used in cell-line-based drug synergy prediction. We compared our model to five baselines used in the drug synergy prediction task: DeepDDS,^19^ AuDNNsynergy,^22^ PRODeepSyn^23^ and DeepSynergy.^20^ In this particular setting, we did not evaluate GraphSynergy due to its requirement for all of the drugs to have known drug target genes. However, we could only collect drug targets for 13 drugs using existing DrugBank and GDSC dataset. We trained our model for 10,000 iterations with a batch size of 64. We set the warm-up steps of the polynomial decay learning scheduler to 2,000. We evaluated the correlations between the predicted volume changes and the observed volume changes using the Spearman correlations coefficients and Pearson correlation coefficients.

### DrugBank and TwoSIDES dataset processing

The drug-drug interaction dataset was collected from the DrugBank and TwoSIDES dataset. We obtained 1,706 drugs with 86 interaction types from the DrugBank dataset and 645 drugs with 963 interaction types. In the raw data, each sample can also be represented as a triplet (drug A, drug B, interaction c), which means the drug A and drug B have an interaction type c. All the triplets appearing in the raw dataset were treated as positive samples. We additionally generated samples that were not included in the original dataset as negative samples. These negative samples were created to match the quantity of the positive samples. We evaluated the vanilla cross validation, one new drug in each test pair and split by combination setting for the DrugBank dataset. The one new drug in each test pair setting means that each combination in the validation set has one drug that is never seen in the training set. The split by combination setting means that all validation combinations never appeared in the training set. We followed the partition procedure in R^2^-DDI^88^ and GMPNN-CS^89^ to split the dataset.

### Comparison approaches for drug-drug interaction predictions

We compared Pisces to six different approaches on drug-drug interaction prediction tasks. **R**^**2**^**-DDI** is a recently proposed DDI prediction approach by refining interaction type features to improve the prediction accuracy. R^2^-DDI simultaneously learned embeddings for different interaction types and drug pairs. The probability of an interaction type between a pair of drugs is calculated using dot product between their embeddings. **GMPNN-CS** utilized a gated message passing mechanism to capture chemical substructure information at different sizes. The final prediction was calculated using the pairs between the learned substructures. **SSI-DDI** utilized graph attention layers to extract substructure interactions from molecular graphs. The final predictions were calculated by aggregating the interaction scores between substructures. **MR-GNN** utilized the multi-resolution based neural networks to capture the node features of molecular graphs and used these features for drug-drug interaction prediction. **MHCADDI** predicts side effects between drugs based on the message passing and co-attention mechanisms. **GAT-DDI** directly used graph attentions for predicting drug-drug interactions. We adopted experimental results of these approaches reported in R^2^-DDI and GMPNN-CS. These results come from validation set accuracy on the DrugBank and TwoSIDES dataset. MHCADDI did not provide their accuracy and AUPRC on the TwoSIDES dataset. The reported AUROC of MHCADDI on TwoSIDES dataset was 0.882, much lower than our method (an AUROC of 0.914), therefore we did not make comparisons in **Fig. 5d-f**.

### Implementation details for drug-drug interaction prediction task

In the DDI prediction task, we aim to predict which type of interaction is associated with one pair of drugs. Here we formulated it as *k*_*c*_ binary classification tasks, where *k*_*c*_ refers to the number of DDI types. We queried the model with an input of triplet (*d*_*a*_, *d*_*b*_, *c*)_*i*_. The output 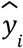 denotes the probability of the input drug pair (*d*_*a*_, *d*_*b*_) has this interaction *c*. To model the interaction type features in each triplet, we assigned a learnable embedding 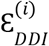 for each interaction type *c* as the additional feature embeddings ***c***. In the augmentor for DDI prediction, the input of the augmentor module only contains a pair of drugs. We first concatenated the drug pair embeddings: 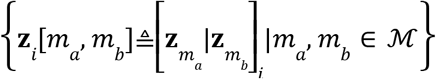. The augmentor transformation layer was used to embed the drug pairs: 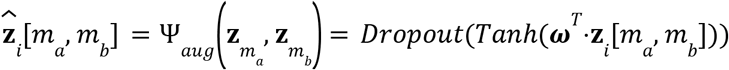 where 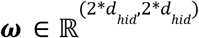. We modified the classification head inspired by R^2^-DDI.^88^ Remember that here we transformed DDI predictions into *k*_*c*_ binary classification tasks and denoted the learnable embeddings of possible interaction types as ***c***. We first embedded the interaction type features 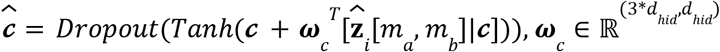. Then we output the predicted probability using a bilinear model: 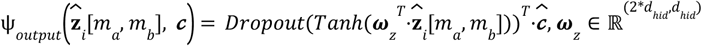.

In the vanilla cross validation setting on both DrugBank dataset and TwoSIDES dataset, we used a 12 layers DeeperGCN and Transformer model. We set the hidden dimension size to 384 and 768 for DeeperGCN and Transformer respectively. The output dimension of the transformation layer *d*_*hid*_ was 512. In the one new drug in each test pair and two new drugs in each test pair setting on DrugBank, we used a 6 layers DeeperGCN with a hidden dimension of 384 and a 6 layers Transformer with the hidden dimension of 512. We set *d*_*hid*_ to 512. {*MPL*_θ[*m*]_ |*m* ∈ ℳ} are one-depth linear layers. We trained the model with a learning rate of 1 × 10^−4^ for 50,000 iterations. We chose the polynomial decay scheduler for the learning rate with 4000 warm-up steps. We ran the grid-search for the dropout rate within [0. 1, 0. 2, 0. 3, 0. 4, 0. 5] and find the optimal dropout rate of 0.1 for vanilla cross validation setting, 0.3 for one new drug in each test pair setting and 0.5 for split by combination setting. The training batch size was set to 128.

We also used predictions from Pisces to construct a DDI network. We first selected ten representative biomedical related drug classes from the drug ontology. Then we obtained 834 drugs, with each of them belonging to at least one of the selected drug classes. We narrowed down our scope on predicting 50 most frequent drug-drug interaction types in DrugBank and collected 868,400 predictions. Next, by filtering out the drug-drug interactions with a predicted probability less than 0.9 and choosing 100 most confident interactions for each drug, we collected 27,510 interactions. We finally applied Fisher’s exact test to identify interaction types that were significantly associated with a pair of classes. Based on these identified interaction types and corresponding drug classes, we were able to generate a DDI network.

## Data Availability

The drug synergy prediction on cell line dataset, drug response prediction on xenografts dataset and the drug-drug interaction dataset are available at https://figshare.com/articles/dataset/Pisces_dataset/23272049.

## Code Availability

Pisces code is available at https://github.com/HanwenXuTHU/Pisces.

