## Supplementary Figure 1, Supplementary Figure 2-4 for "Pisces: A multi-modal data augmentation approach for drug combination synergy prediction"

### Supplementary Infomation

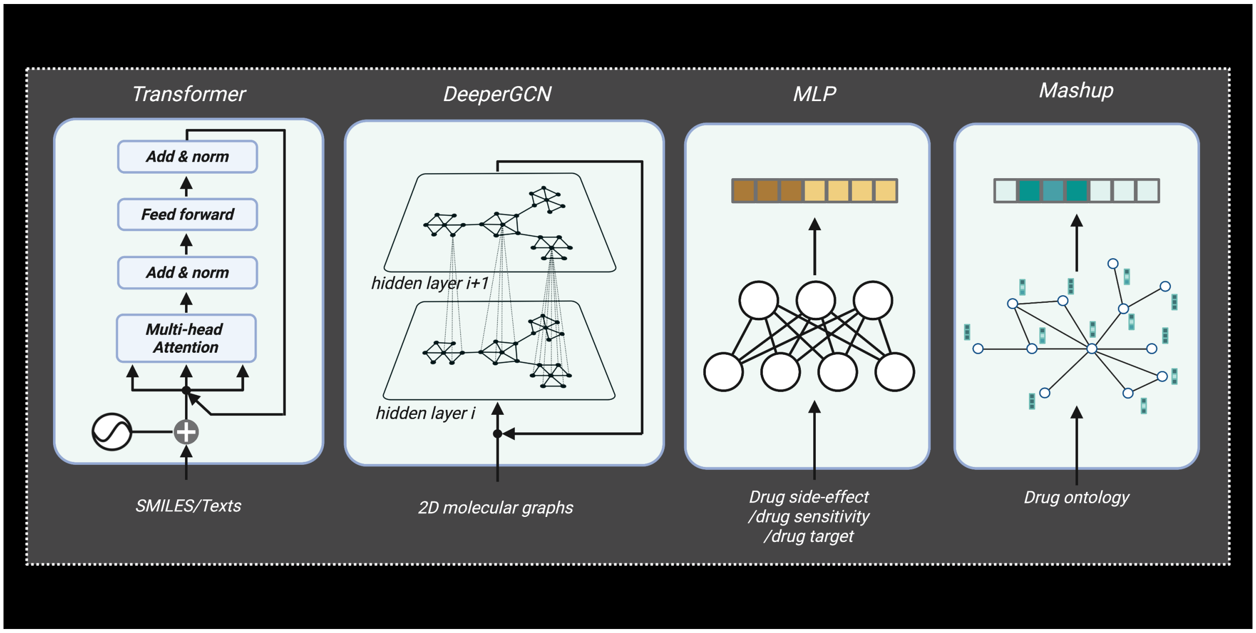


**Supplementary Fig. 1 Model architectures of encoders for different modalities.** We used Transformer architectures for SMILES and textual descriptions, DeeperGCN for molecular graphs (including encoders aligned with 3D geometric views). We used MLP layers for drug sensitivity, drug side effects and drug targets. We finally used Mashup to extract the drug ontology structures.

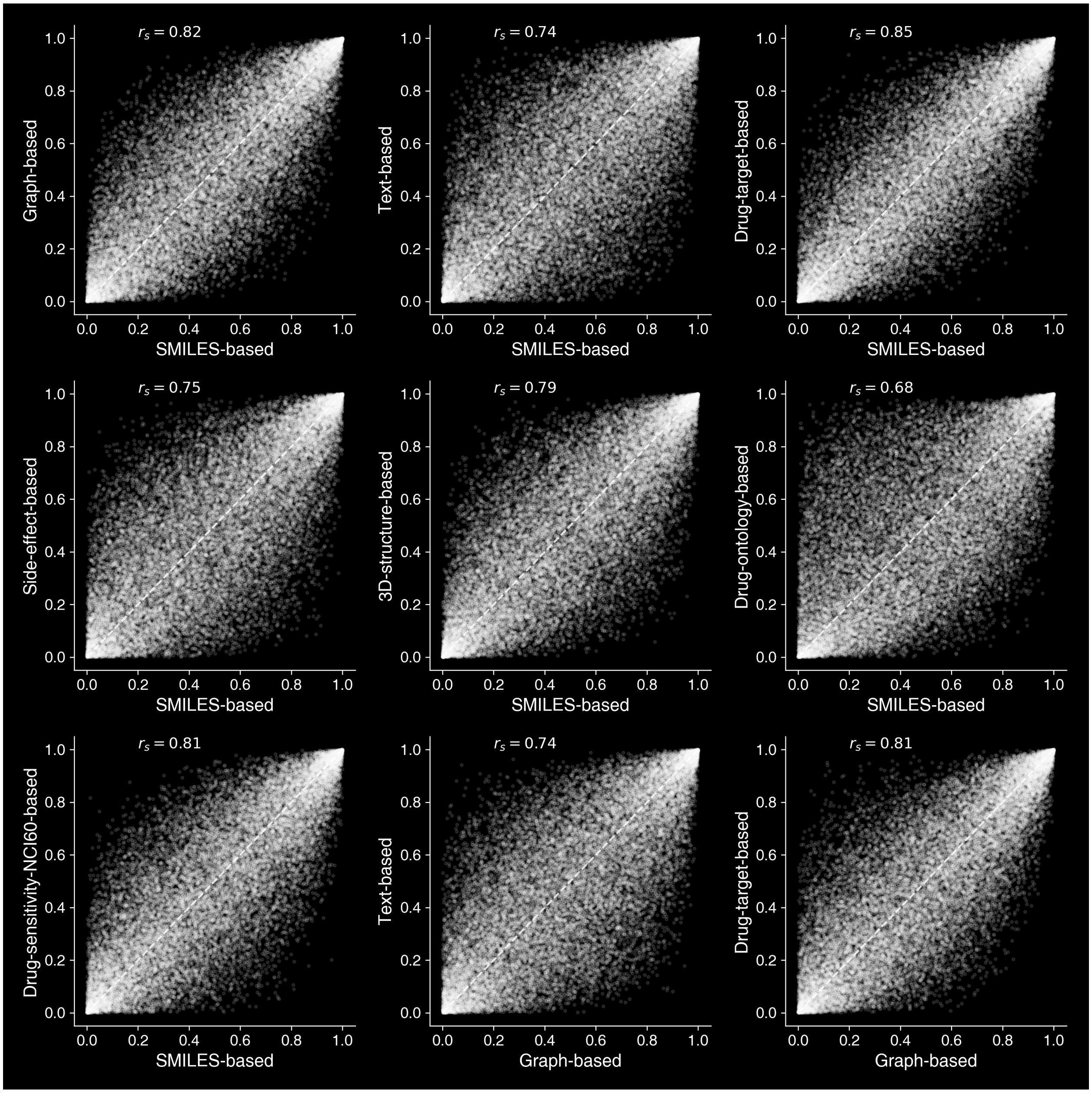


**Supplementary Fig. 2 The agreement among predictions of different modalities.** Scatter plot comparing the prediction scores of using different modalities. We separately performed normalized ranking for predictions based on each modality. The correlations between modalities were measured using Spearman correlation.

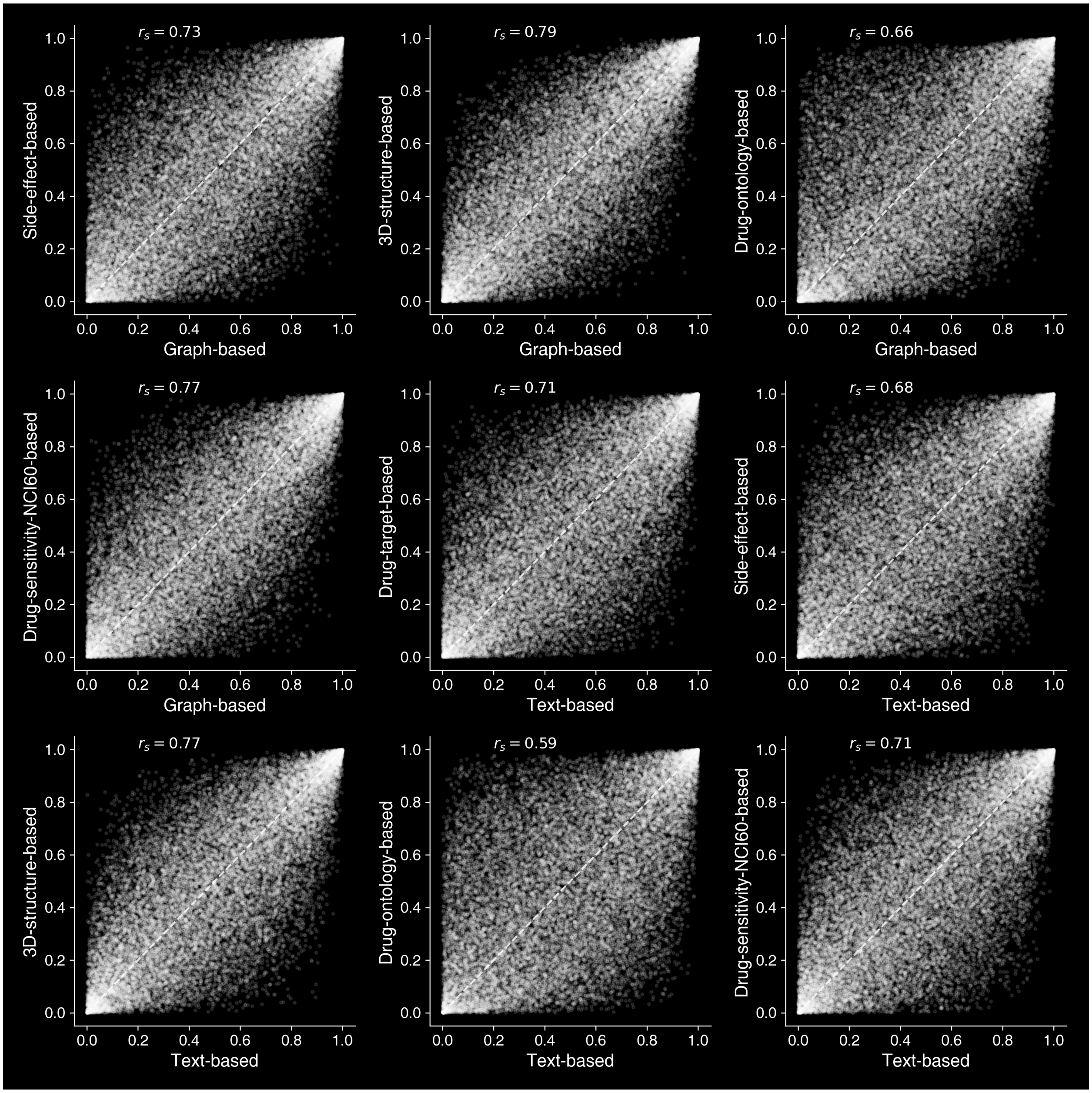


**Supplementary Fig. 3 The agreement among predictions of different modalities.** Scatter plot comparing the prediction scores of using different modalities. We separately performed normalized ranking for predictions based on each modality. The correlations between modalities were measured using Spearman correlation.

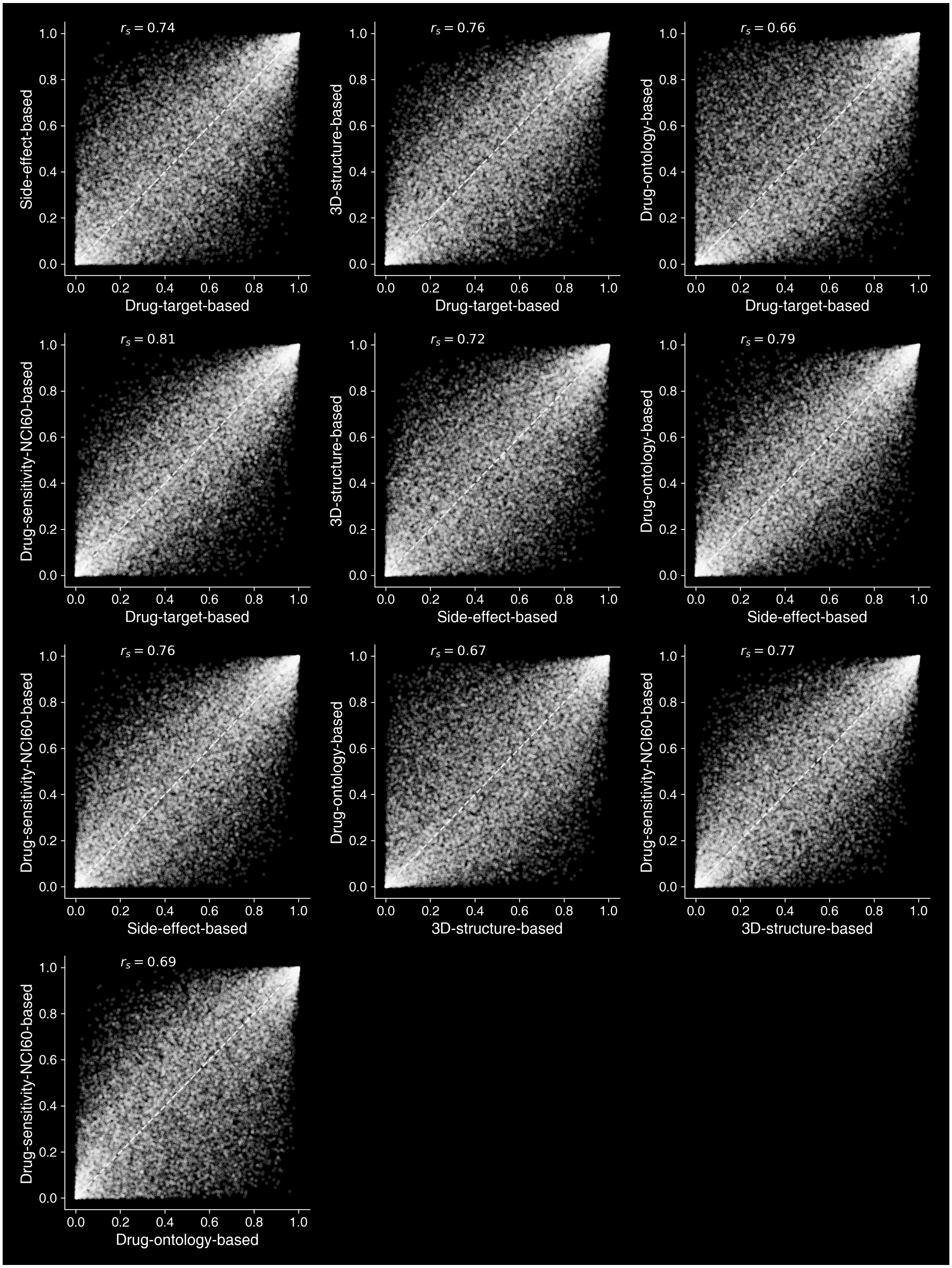


**Supplementary Fig. 4 The agreement among predictions of different modalities.** Scatter plot comparing the prediction scores of using different modalities. We separately performed normalized ranking for predictions based on each modality. The correlations between modalities were measured using Spearman correlation.

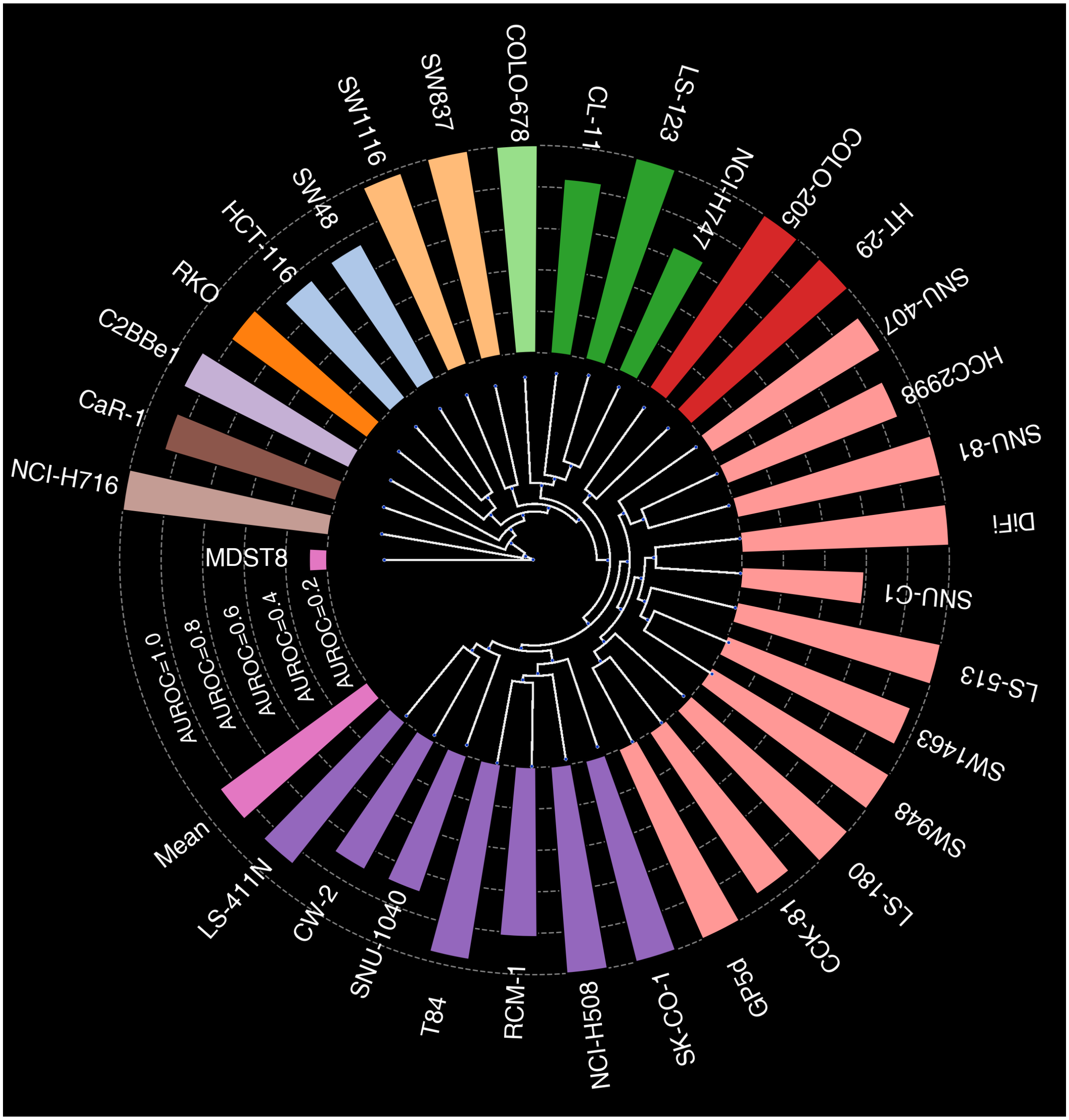


**Supplementary Fig. 5 Three-drug combination prediction in terms of AUROC.** Circular bar plots showing Pisces' prediction performance in terms of AUROC on three-drug combination prediction when trained only on two-drug and single-drug data. AUROC are only shown stratified by cell line features. The circular dendrogram shows the hierarchical clustering of cell lines using gene expression levels. The branch height represents the distances between two cell line clusters.

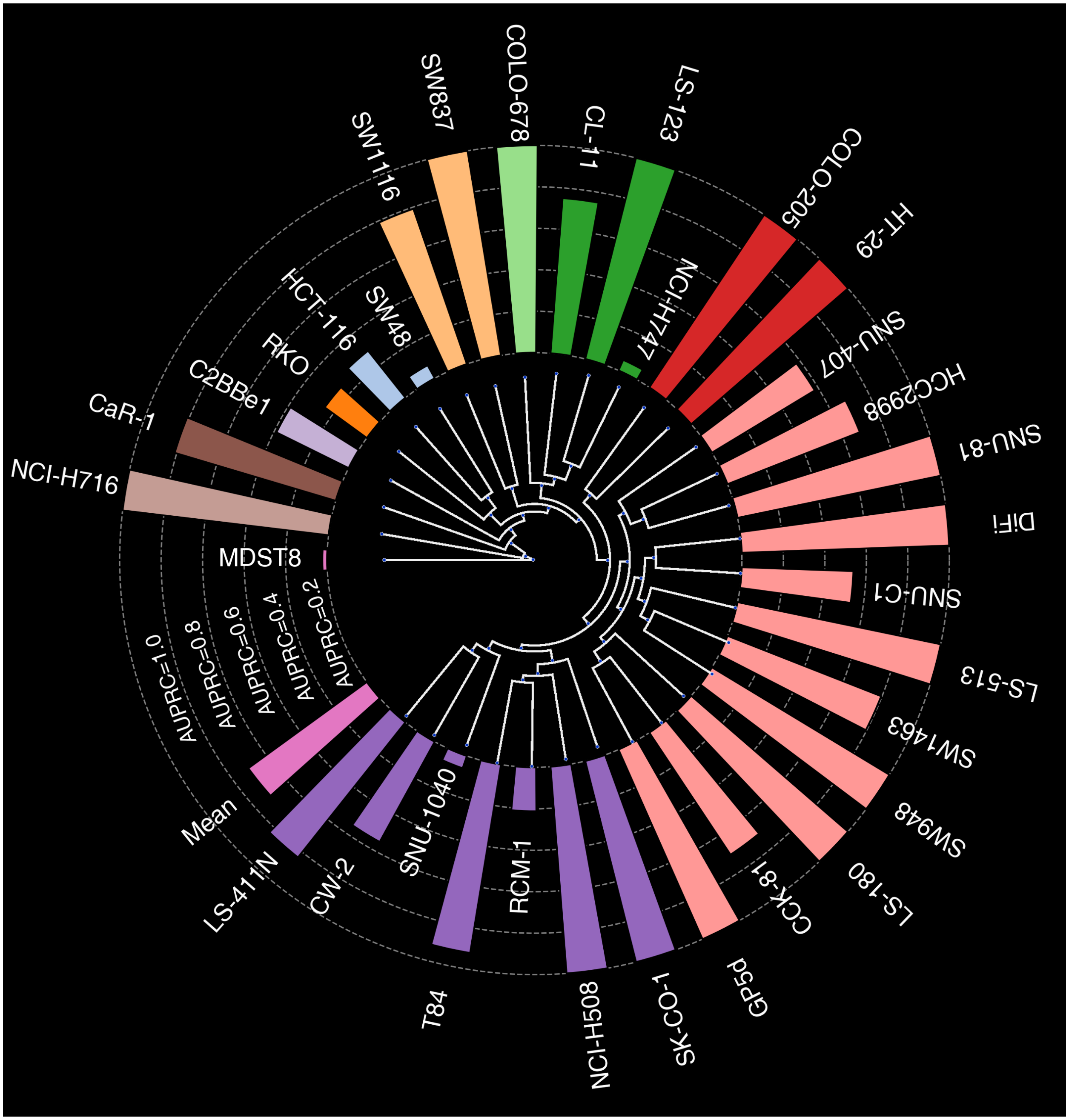


**Supplementary Fig. 6 Three-drug combination prediction in terms of AUPRC.** Circular bar plots showing Pisces' prediction performance in terms of AUPRC on three-drug combination prediction when trained only on two-drug and single-drug data. AUPRC and AUPRC are only shown stratified by cell line features. The circular dendrogram shows the hierarchical clustering of cell lines using gene expression levels. The branch height represents the distances between two cell line clusters.

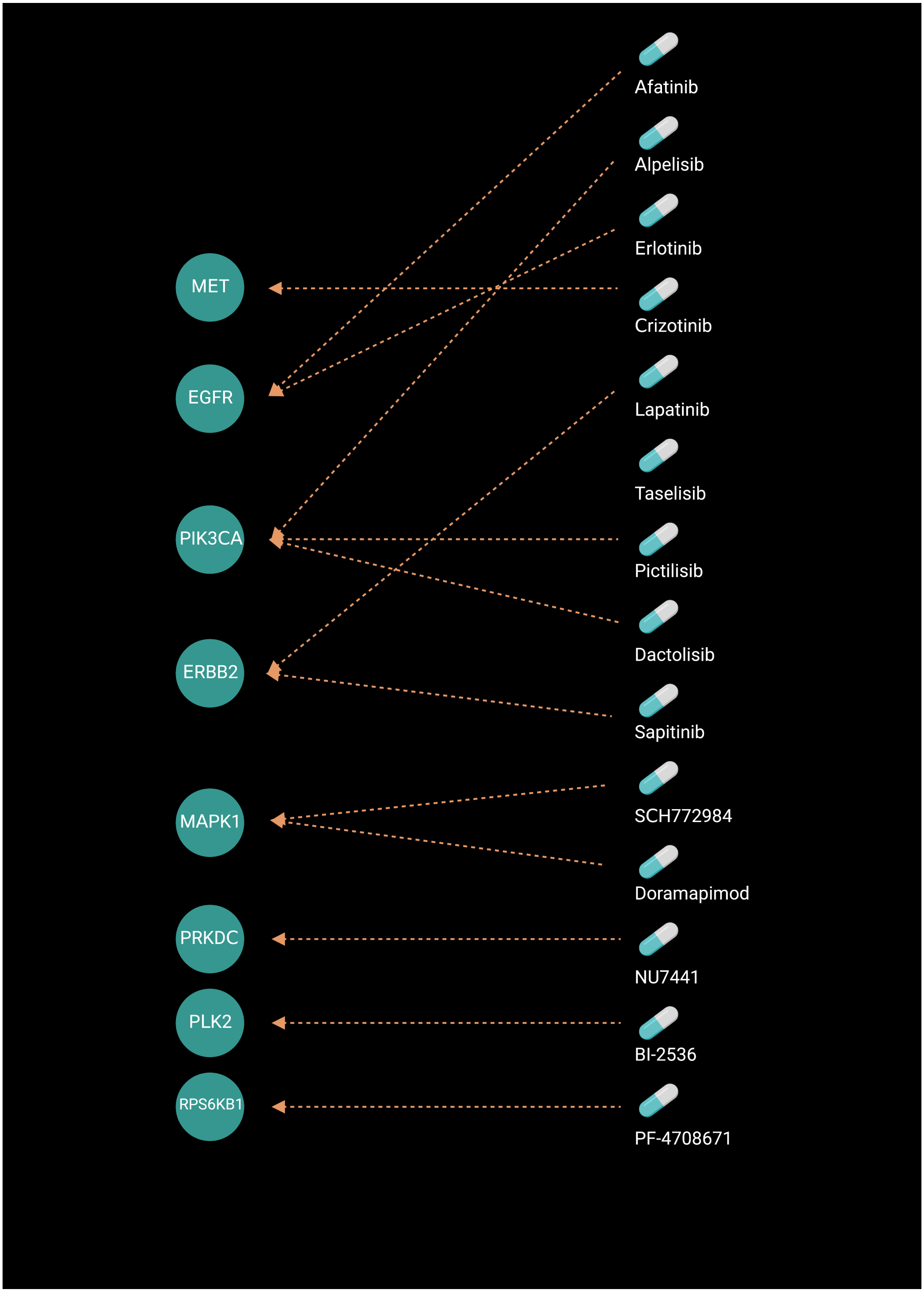


**Supplementary Fig. 7 Bipartite graph of drugs and target genes.** Bipartite graph showing 14 cancer drugs and 8 genes that can be inhibited by one of these drugs. Each edge links one drug and the gene that can be inhibited by this drug.

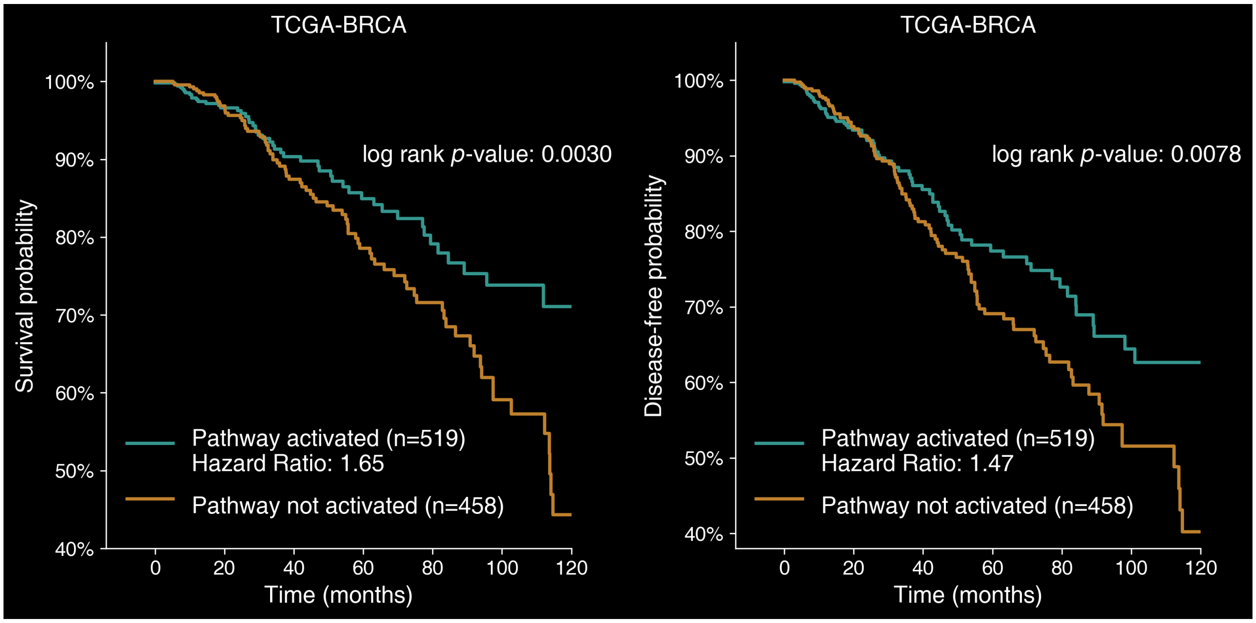


**Supplementary Fig. 8 Survival analysis on patients from TCGA dataset.** Survival plots showing the significantly different overall survival (left) and disease-free survival (right) between two groups of TCGA-BRCA patients. These two groups were clustered based on whether the BRCA drug-sensitive pathway was activated using the gene expression.

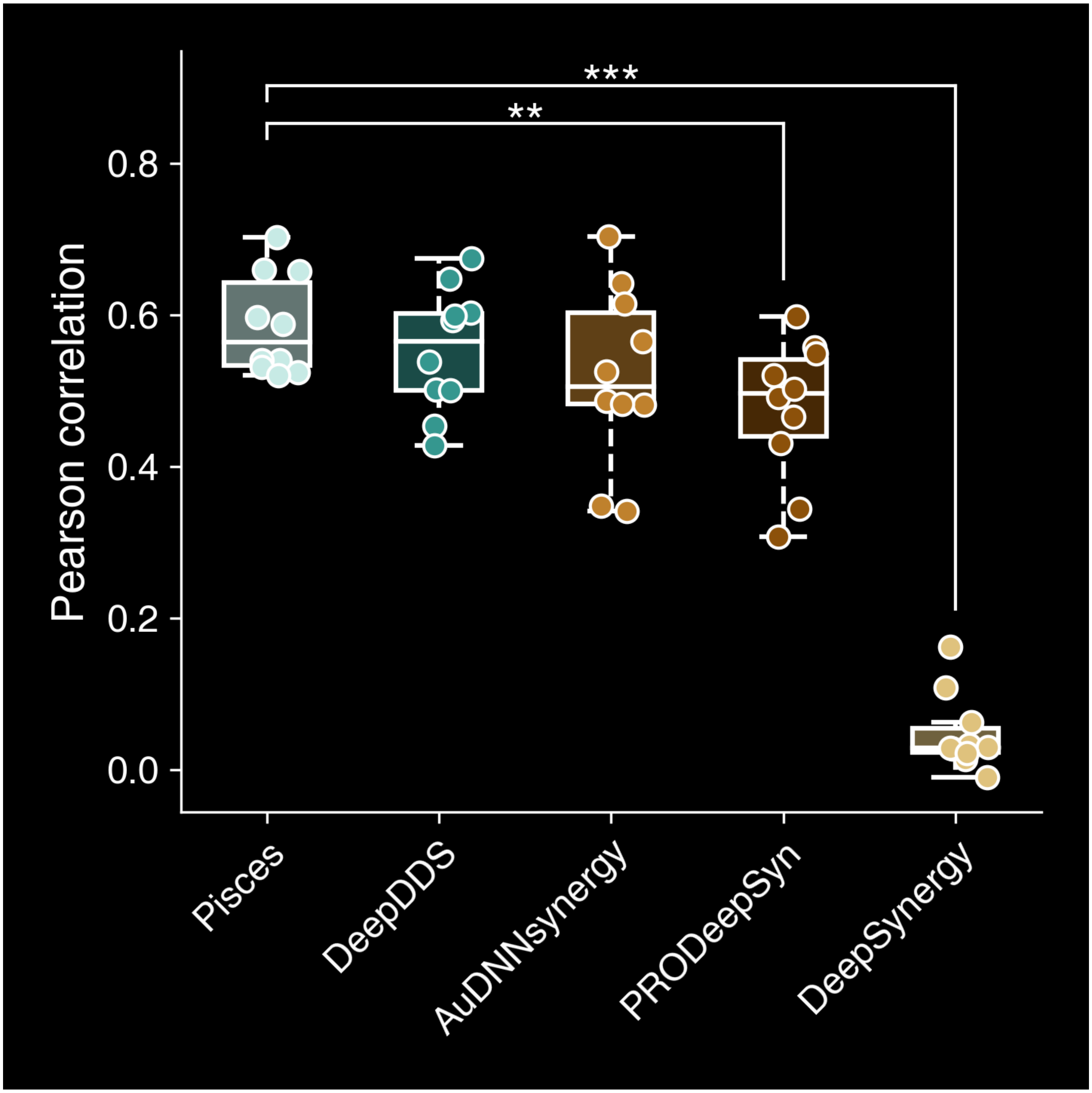


**Supplementary Fig. 9 Tumor change prediction in terms of Pearson correlation.** Box plot comparing the drug synergy prediction on xenografts using Pearson correlation. The * indicates that Pisces outperforms the next-best-performing model in the metric, with significance levels of t-test $p$-value $< 5 \times 10^{-2}$ for *, t-test $p$-value $< 1 \times 10^{-2}$ for **, and t-test $p$-value $< 5 \times 10^{-3}$ for ***. All t-tests are one-sided.

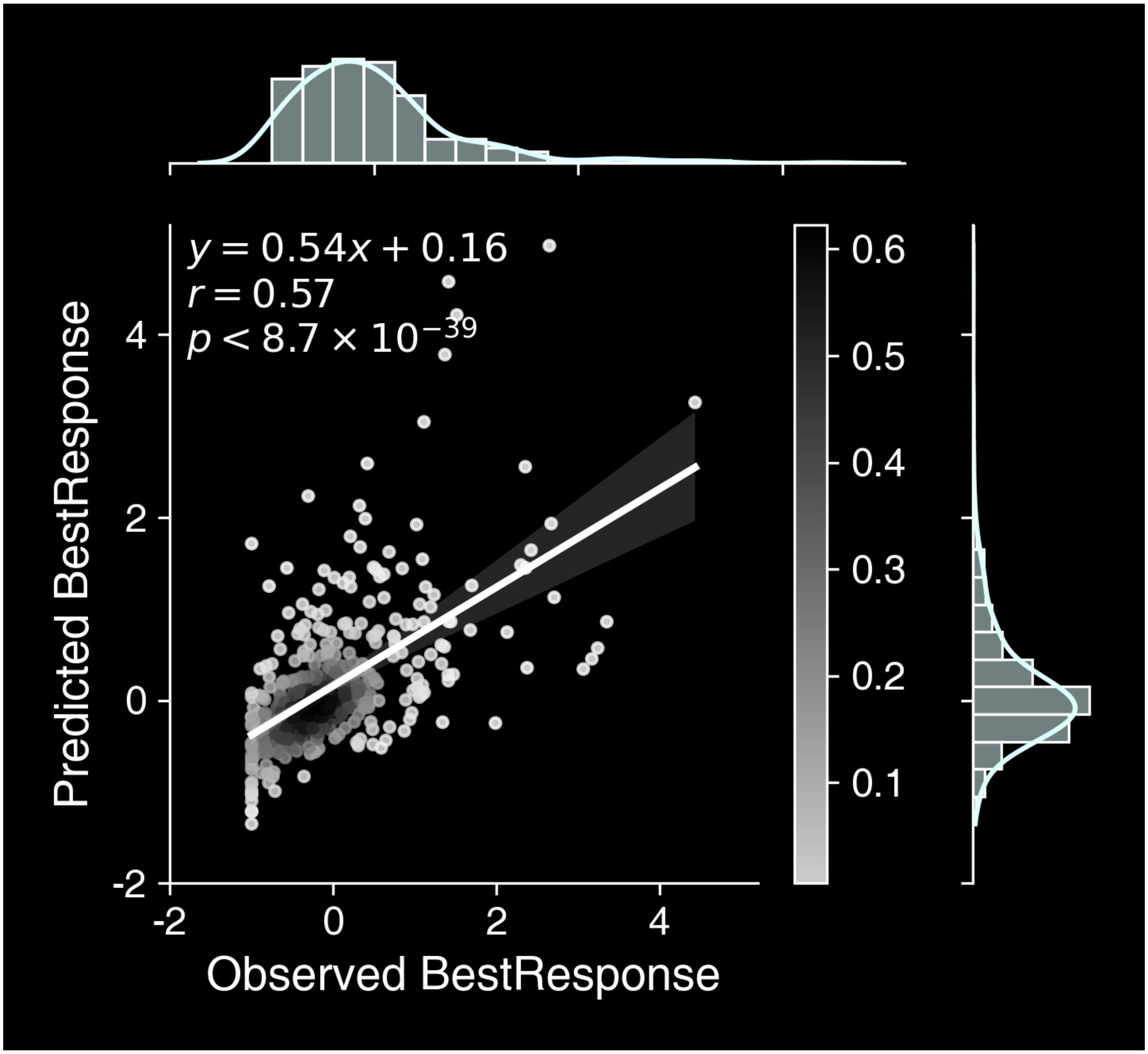


**Supplementary Fig. 10 Scatter plot of tumor change prediction.** Scatter plots comparing the predicted and observed minimum %tumor changes after 10 days at holdout triplets. Bar plots show the frequency.

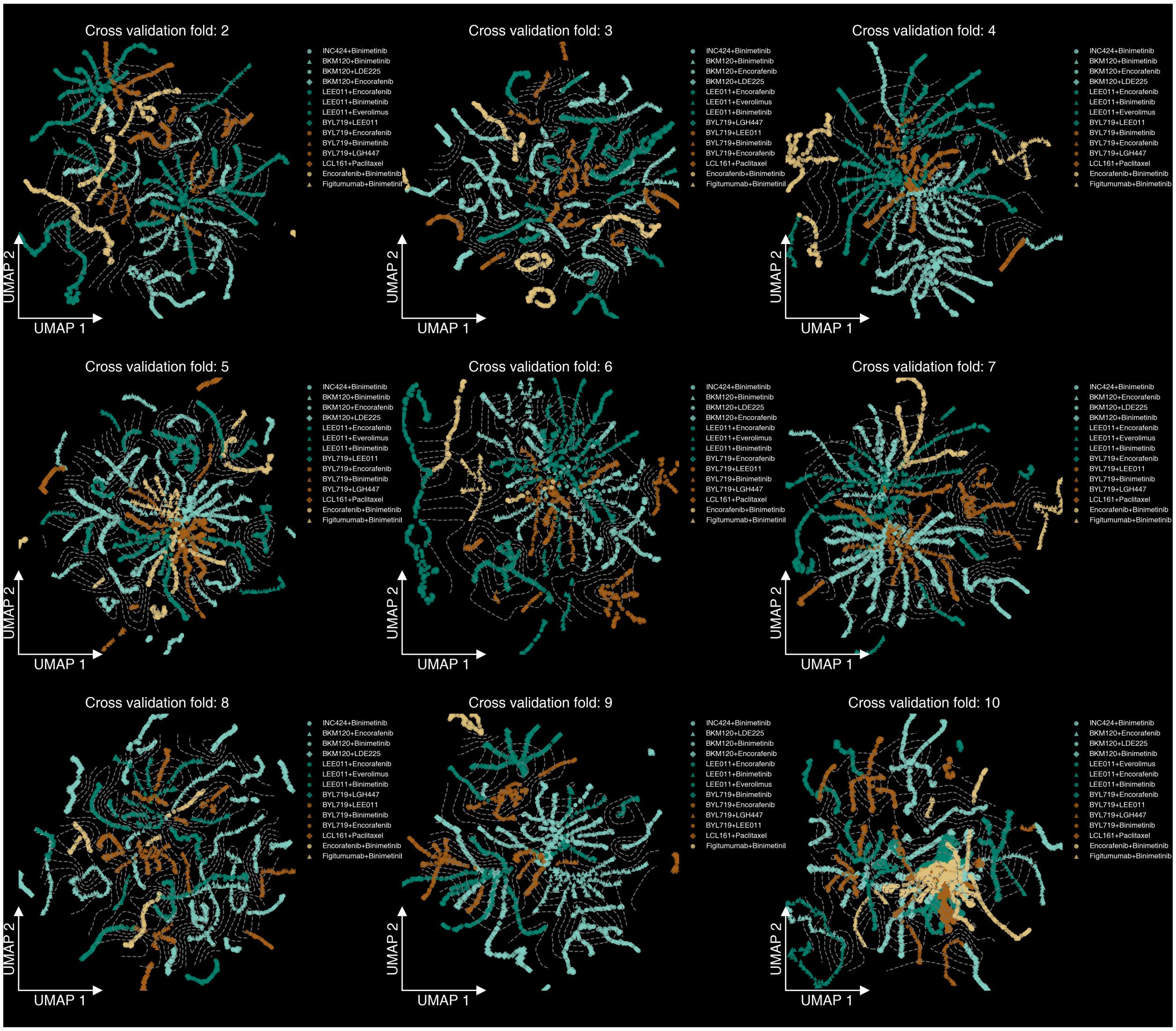


**Supplementary Fig. 11 UMAP plot of test triplets at different time points.** UMAP plot showing the embedding of test triplets at different time points from 9 other folds in the cross validation. Each triplet is a pair of drugs and a xenograft model. Nodes are colored and marked by the drug pair.

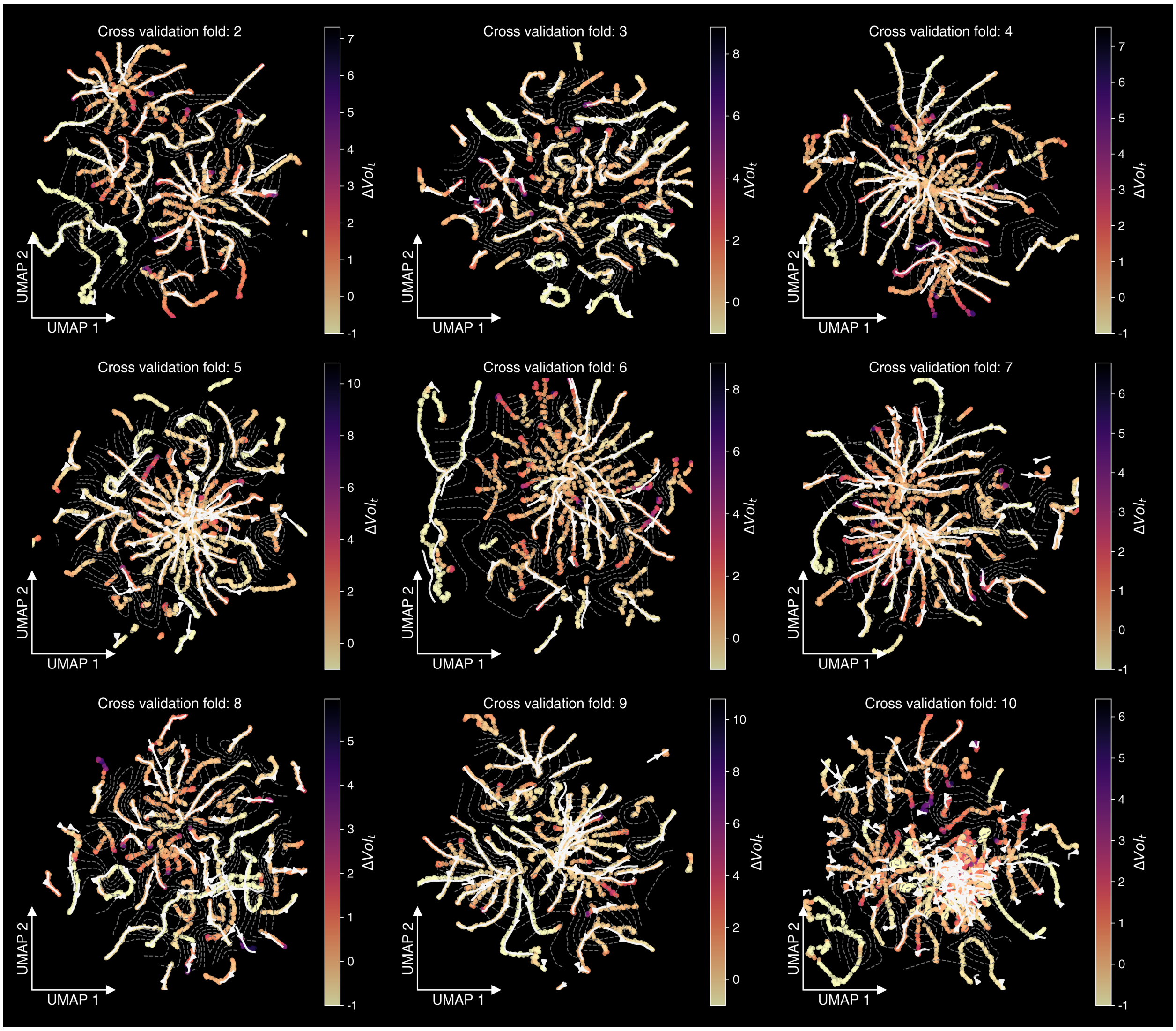


**Supplementary Fig. 12 UMAP plot of one drug pair on different xenografts.** UMAP plot showing the embedding of test triplets at different time points from 9 other folds in the cross validation. Each triplet is a pair of drugs and a xenograft model. The nodes are colored and marked by the tumor volume change. The contours connect tumors that have the same time point. The arrows are from early time points to later time points.

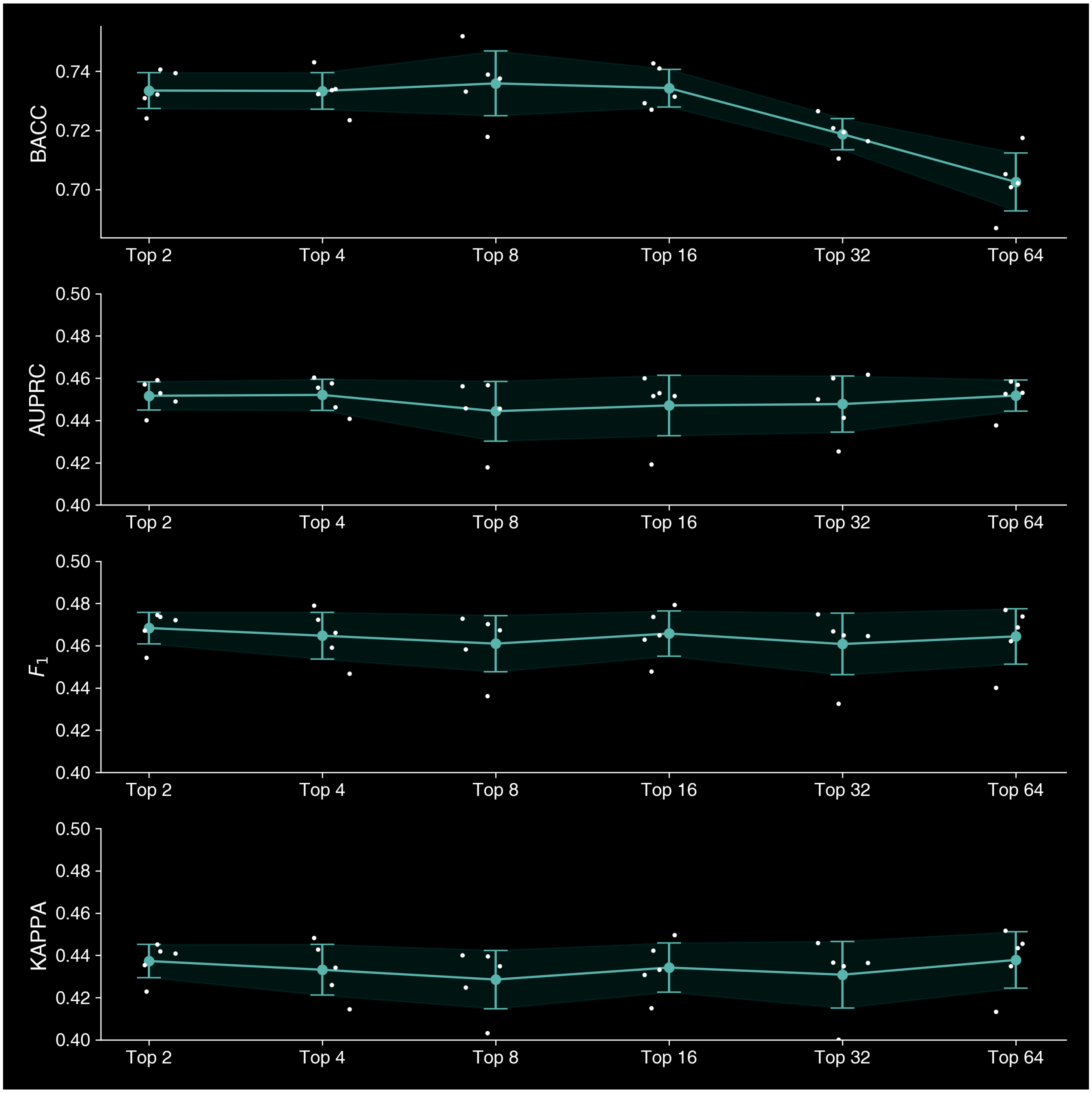


**Supplementary Fig. 13 Ablation studies on various top k.** UMAP plot showing the performance changes by aggregating various top k predictions. The x axis show the aggregation of top 2, 4, 8, 16, 32 and all 64 predictions. We investigated on the GDSC-Combo dataset in the vanilla cross validation setting. Performances were measured in terms of BACC, AUPRC, $F_1$ and KAPPA. Error bar represents the standard deviation across 5 fold cross validation. Each white dot represents on one fold.

| Drug A Class | Drug B Class | Drug-drug interactions | *p*-value | Number of interactions |
|---|---|---|---|---|
| phenols | homopolycyclic compound | Increase the anticholinergic activities | $5.63\times10^{-9}$ | 43 |
| zwitterion | organofluorine compound | Increase the risk or severity of QTc prolongation | $2.40\times10^{-4}$ | 31 |
| organochlorine compound | phenols | Decrease the vasoconstricting activities | $1.12\times10^{-10}$ | 26 |
| zwitterion | organochlorine compound | Decrease the diuretic activities | $1.30\times10^{-7}$,26 | |
| organochlorine compound | homopolycyclic compound | Increase the constipating activities | $4.80\times10^{-7}$ | 22 |
| organic fundamental parent | homopolycyclic compound | Decrease the sedative activities | $1.54\times10^{-9}$,21 | |
| organochlorine compound | hydroxy steroid | Increase the thrombogenic activities | $3.59\times10^{-10}$ | 18 |
| organofluorine compound | phenols | Decrease the vasoconstricting activities | $3.13\times10^{-11}$ | 17 |
| phenols | organic fundamental parent | Increase the anticholinergic activities | $6.03\times10^{-7}$ | 17 |
| homopolycyclic compound | phenols | Increase the thrombogenic activities | $8.91\times10^{-5}$ | 16 |
| organofluorine compound | homopolycyclic compound | Increase the anticholinergic activities | $2.64\times10^{-5}$ | 14 |
| organic fundamental parent | organofluorine compound | Increase the thrombogenic activities | $2.93\times10^{-4}$ | 14 |
| aromatic amine | homopolycyclic compound | Decrease the sedative activities | $2.40\times10^{-7}$ | 12 |
| sulfonic acid derivative | organochlorine compound | Decrease the diuretic activities | $4.21\times10^{-3}$ | 9 |
| sulfonic acid derivative | hydroxy steroid | Increase the thrombogenic activities | $1.52\times10^{-6}$ | 9 |
| aromatic amine | hydroxy steroid | Increase the thrombogenic activities | $1.05\times10^{-5}$ | 6 |
| olefinic compound | hydroxy steroid | Increase the thrombogenic activities | $4.61\times10^{-3}$ | 5 |
| organofluorine compound | organic fundamental parent | Decrease the vasoconstricting activities | $5.83\times10^{-5}$ | 4 |
| olefinic compound | phenols | Increase the anticholinergic activities | $2.99\times10^{-4}$ | 3 |
| homopolycyclic compound | hydroxy steroid | Increase the thrombogenic activities | $8.17\times10^{-3}$ | 3 |
| sulfonic acid derivative | organic fundamental parent | Increase the risk or severity of QTc prolongation | $1.75\times10^{-4}$ | 3 |

**Supplementary Table 1** Table showing drug-drug interactions significantly occurred between two drug classes. The first three columns indicate two drug classes and the associated interaction type . The *p*-value represented the Fisher's exact test results. The number of interactions means how many interactions found between drugs from these two classes.
